# Chronic Stress Alters Dorsal Bed Nucleus of Stria Terminalis Synaptic Neurotransmission in a Dravet Syndrome Mouse Model

**DOI:** 10.64898/2026.05.19.723288

**Authors:** Eunyoung Hong, Emily Y Xu, Jackson G Murray, Jiarui Qin, Sarah M Mulloy, Yana Van den Abeele, Lavanya Dhavala, Jayne A Miner, Gabi R Barrocas, Betzy M Martinez, Alyssa A Mitchell, Flavia Villa, William P Nobis

**Affiliations:** Department of Neurology, Vanderbilt University Medical Center, Nashville, Tennessee USA; Vanderbilt Brain Institute, Vanderbilt University, Nashville, Tennessee USA; Vanderbilt University, Nashville, Tennessee USA

## Abstract

Stress is a commonly reported seizure precipitant and may contribute to the development of psychiatric comorbidities in epilepsy, yet how chronic stress interacts with epileptic circuits remains poorly understood. We investigated the impact of chronic restraint stress on physiological, behavioral, and synaptic outcomes in a mouse model of Dravet syndrome, specifically corticotropin-releasing factor (CRF) neurons in the bed nucleus of the stria terminalis (BNST), a stress-responsive region implicated in epilepsy patients. Chronic restraint stress produced divergent hypothalamic-pituitary-adrenal axis responses, with stressed Dravet syndrome mice exhibiting elevated corticosterone, increased mortality in females, and increased locomotion and anxiety-like behavior. Ex vivo electrophysiological recordings revealed that chronic stress increased spontaneous excitatory event frequency onto BNST CRF neurons in both genotypes and selectively increased sEPSC and sIPSC amplitude in Dravet syndrome mice. Evoked recordings demonstrated genotype-specific effects of stress on glutamatergic transmission in CRF neurons of the DS group. This suggests greater stress-dependent remodeling of spontaneous and evoked synaptic activity in DS. These findings suggest chronic stress may worsen physiological and behavioral outcomes in Dravet syndrome and promote specific maladaptive alterations in BNST CRF circuitry. More broadly, these results suggest that stress interacts with seizure vulnerability and potentially contributes to neuropsychiatric comorbidities and epilepsy.

## Introduction

Epilepsy is a prevalent neurological disorder that affects approximately 50 million people worldwide (Chen et al. 2023). It is characterized by unpredictable seizures, which can significantly disrupt basic lifestyle routines (England et al. 2012b, 2012a). Patients with epilepsy present a disproportionately high risk for psychiatric comorbidities, with at least 1 in 3 patients having a psychiatric diagnosis (Mula et al. 2021; Gurgu et al. 2021). Stress is reported as a seizure precipitant in up to 85% of people with epilepsy, and this relationship is likely bidirectional, as seizures may elevate physiological and psychological stress which in turn may reduce seizure threshold (McKee and Privitera 2017; Lang et al. 2018; Catalán-Aguilar et al. 2025; Lathers and Schraeder 2006). Thus, better understanding how seizures interact with the neural substrates underlying stress may inform treatment approaches and improve quality of life (Gandy et al. 2021; Espinosa-Garcia et al. 2021).

The bed nucleus of the stria terminalis (BNST) is a region within the extended amygdala that plays a critical role in sustained stress and anxiety-like behavior (Daniel and Rainnie 2016; L. Tran et al. 2014; Lebow and Chen 2016; S.-Y. Kim et al. 2013). In preclinical models, our lab has shown that seizures activate the BNST in both the DBA/1J audiogenic seizure model and a transgenic mouse model of Dravet syndrome, and viral inhibition of the dorsolateral BNST (dlBNST) mitigates seizure-induced respiratory arrest (Xia et al. 2022; Yan et al. 2021). Clinically, patients with temporal lobe epilepsy (TLE) exhibit reduced structural and effective connectivity between the BNST and both cortical regions and subcortical arousal centers (Reda et al. 2025). Additionally, the BNST is enlarged in TLE patients, with even greater enlargement observed in those with comorbid depression (Dhaher et al. 2022). These findings suggest that epilepsy may drive synaptic changes within the BNST that contribute to the development of comorbid mood disorders.

Dravet syndrome (DS) is a genetic epileptic encephalopathy primarily caused by a loss-of-function mutation in the Scn1a gene (Yan et al. 2021; Ricobaraza et al. 2019; Kaneko et al. 2022; Prentice et al. 2026). Beyond early-life febrile seizures, individuals with DS carry a high risk of sudden unexpected death in epilepsy (SUDEP) and a prominent neuropsychiatric phenotype including developmental delay, social impairment, and psychiatric comorbidities (Ricobaraza et al. 2019). Individuals with DS have more seizure precipitants and respond more severely to physical stress compared to other childhood epilepsies (Verbeek et al. 2015). Rodent models reliably recapitulate this phenotype, making them a translatable model for mechanistic investigation (Valassina et al. 2022; Sawyer et al. 2016). The BNST is particularly relevant in this context, as it contains one of the highest concentrations of stress-activated corticotropin-releasing factor (CRF) neurons (Maita et al. 2022; Dabrowska et al. 2016; Partridge et al. 2016). Furthermore, we have previously shown that DS mice exhibit altered excitatory synaptic transmission within the BNST (Yan et al. 2021).

Little investigation has been done on the interplay between stress and epilepsy, particularly in epileptic syndromes such as DS where stressors may more readily precipitate seizures due to altered stress responses. In this study, we addressed this gap by examining the impact of chronic restraint stress on behavior, mortality, and the synaptic transmission in the BNST of DS mice.

## Results

To investigate if the physiological response to chronic stress is different in DS mice, we monitored daily weight, serum CORT levels, and mortality (**Fig. 1A**). Both stressed wild-type (WT) and DS mice gained significantly less weight than their respective controls across the 8 day restraint period. Chronic stress significantly lowered normalized weight gain in WT mice compared to controls at every time point from day 2 through day 8 (p < 0.0001 for all time points, **Fig. 1B**). Similarly, chronic stress lowered weight gain in DS mice at every time point from day 2 to day 8 (p = 0.0002 for day 2, p < 0.0001 for day 3-8, **Fig. 1C**). There were no differences in weight gain between control WT and control DS mice or between stressed WT and stressed DS mice.

**Figure 1:**
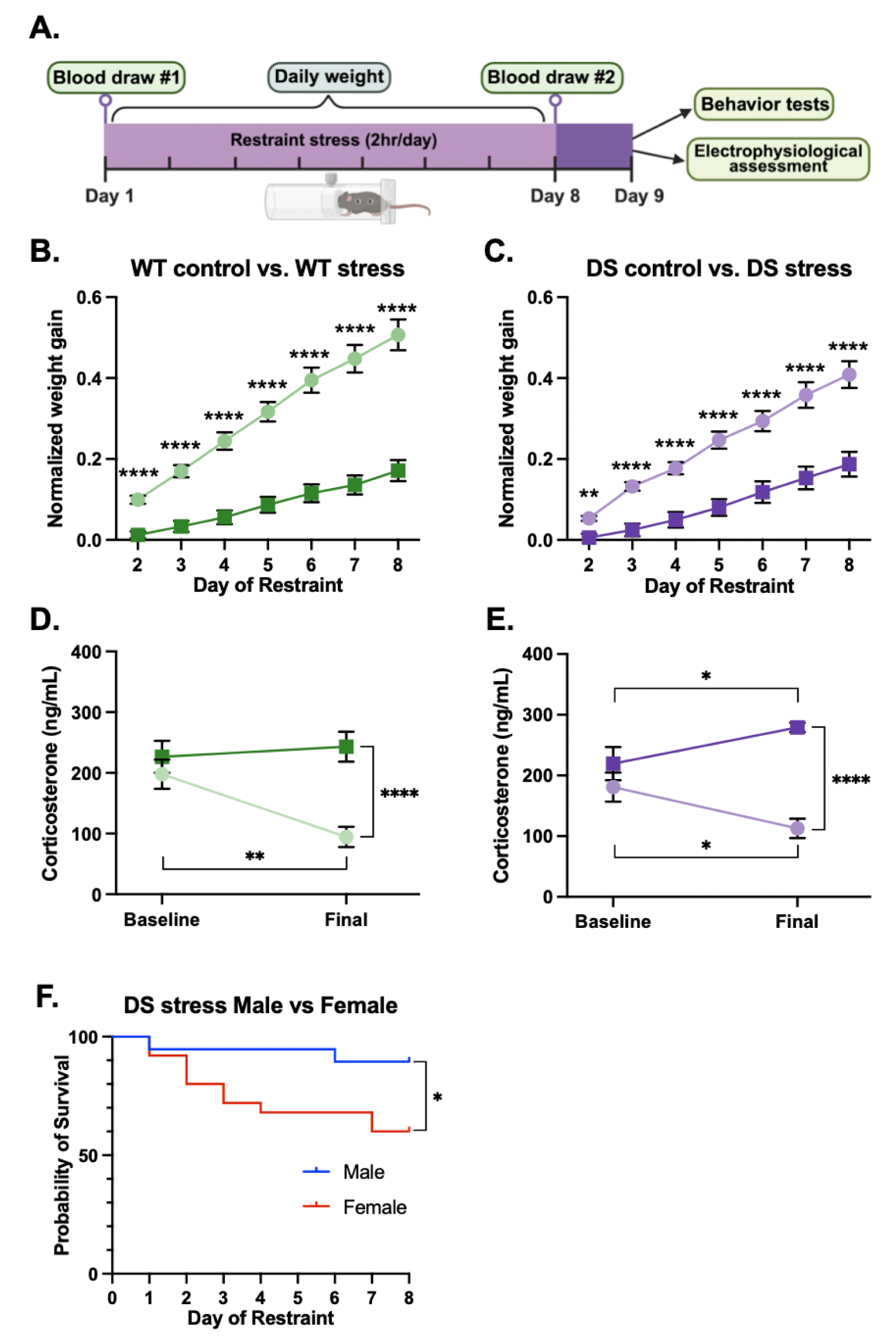
Chronic restraint stress reduces weight gain, elevates corticosterone levels, and increases SUDEP. **(A)** Experimental timeline. Mice underwent 8 days of chronic restraint stress (2 hours/day) beginning on day 1, with daily weight measurements. Tail blood was collected for corticosterone (CORT) measurement following the first restraint session (baseline) and on day 9 (final). Behavior testing and electrophysiological assessments were performed on day 9. Four groups were examined: WT control (light green), WT stress (dark green), DS control (light purple), and DS stress (dark purple). **(B)** Normalized weight gain in WT control (n = 21) and WT stress (n = 24) mice (two-way repeated measures ANOVA, time: F(1.31, 56.33) = 241.7, p < 0.0001; stress: F(1, 43) = 55.76, p < 0.0001; time × stress interaction: F(1.31, 56.33) = 48.29, p < 0.0001; Benjamini-Hochberg FDR post-hoc test; p < 0.0001 at all time points). **(C)** Normalized weight gain in DS control (n = 21) and DS stress (n = 22) mice (two-way repeated measures ANOVA, time: F(1.35, 55.32) = 121.9, p < 0.0001, stress: F(1, 41) = 29.45, p < 0.0001, time × stress interaction: F(1.35, 55.32) = 11.98, p < 0.001). **(D)** Serum CORT levels at baseline and following the final restraint session in WT control (n = 10) and WT stress (n = 10) mice (two-way repeated measures ANOVA; time: F(1, 18) = 3.94, p = 0.063, stress: F(1, 18) = 12.99, p = 0.002, time × stress interaction: F(1, 18) = 7.55, p = 0.013). **(E)** Serum CORT levels at baseline and following the final restraint session in DS control (n = 10) and DS stress (n = 9) mice (two-way repeated measures ANOVA; time: F(1, 17) = 0.057, p = 0.81, stress: F(1, 17) = 20.11, p < 0.001, time × stress interaction: F(1, 17) = 12.97, p = 0.002). **(F)** Kaplan-Meier survival curves for DS stressed male (n = 19) and female (n = 25) mice. Female mice showed significantly reduced survival compared to males (log-rank test: χ²(1) = 4.45, p = 0.035). No deaths were observed in WT mice under any condition. Statistical significance was determined with Benjamini-Hochberg FDR **(B, C)** or Fisher’s LSD **(D, E)** post hoc tests were used. *p < 0.05, **p < 0.01, ***p < 0.001, ****p < 0.0001.

Furthermore, to confirm that our restraint protocol engaged the HPA axis throughout the 8 day period, we measured CORT levels following the first and final restraint session. Following the first restraint, CORT levels did not differ between control and stressed mice in either genotype. On the final day, control mice in both genotypes showed a significant reduction in CORT levels from baseline. Notably, chronic stress evoked a stable CORT response in the WT mice (p = 0.60, **Fig. 1D**), whereas DS mice showed a significant increase from baseline (p = 0.033, **Fig. 1E**).

Finally, given that Dravet syndrome is associated with an elevated risk of SUDEP, we monitored mortality across the 8 day restraint period. No deaths were observed in WT mice regardless of condition. In stressed DS mice, female mice exhibited a significantly increased mortality rate compared to males (log-rank test: χ²(1) = 4.45, p = 0.035; **Fig. 1F**). This sex difference was not observed in control DS mice. Moreover, stressed DS females had a higher mortality rate than the nonstressed DS females (log-rank test: χ²(1) = 4.23, p = 0.04), whereas this difference was not observed in the DS males (log-rank test: χ²(1) = 2.06, p = 0.15).

Anxiety-like behavior was assessed using the zero maze (EZM) on day 9 following the last day of stress (**Fig. 2A**, **Fig. 1A)**. Stressed DS mice traveled a significantly greater distance compared to control DS (p < 0.001) and stressed WT mice (p = 0.019, **Fig. 2B**). In comparison, stressed WT mice spent more time in the open arm compared to WT controls (p = 0.004, **Fig. 2C**). Furthermore, we quantified the frequency of head dips as a measure of exploratory risk assessment behavior. Stressed DS mice made significantly more head dips than DS controls (p = 0.003, **Fig. 2D**).

**Figure 2:**
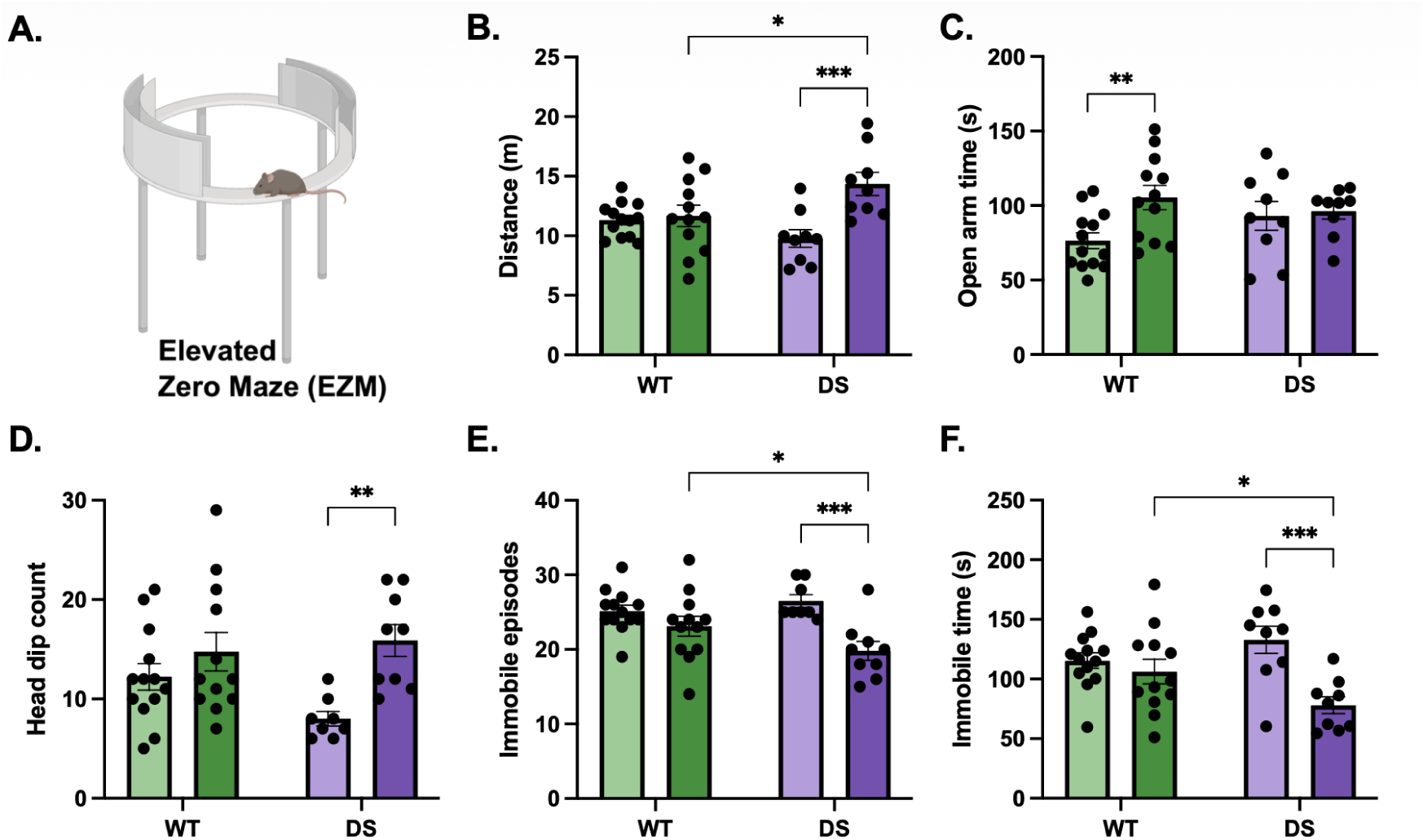
Chronic stress produces distinct anxiety-like behavioral changes in DS mice. **(A)** Schematic of the elevated zero maze (EZM). Mice were tested on day 9 following the final session for WT control (light green), WT stress (dark green), DS control (light purple), and DS stress (dark purple) mice (n=9-13 animals per group). **(B)** Total distance traveled on the EZM (genotype: F(1, 39) = 0.53, p = 0.47; stress: F(1, 39) = 10.53, p = 0.002; genotype × stress interaction: F(1, 39) = 7.70, p = 0.008). **(C)** Time spent in the open arms (genotype: F(1, 39) = 0.26, p = 0.61; stress: F(1, 39) = 4.80, p = 0.035; genotype × stress interaction: F(1, 39) = 3.11, p = 0.086). **(D)** Number of total head dips while positioned on the open arm (genotype: F(1, 38) = 0.93, p = 0.34; stress: F(1, 38) = 10.54, p = 0.002; genotype × stress interaction: F(1, 38) = 2.81, p = 0.10). **(E)** Number of total immobile episodes (genotype: F(1, 38) = 0.75, p = 0.39; stress: F(1,38) = 15.18, p < 0.001; genotype × stress interaction: F(1,3 8) = 4.25, p = 0.046). **(F)** Total time spent immobile (genotype: F(1, 39) = 0.34, p = 0.56; stress: F(1, 39) = 12.40, p = 0.001; genotype × stress interaction: F(1,3 9) = 6.31, p = 0.016). Statistical significance was determined by 2-way ANOVA followed by Fisher’s LSD post hoc test. *p < 0.05, **p < 0.01, ***p < 0.001.

Additionally, stressed DS mice had significantly fewer immobile episodes and decreased total immobile time compared to both DS controls (immobile episodes: p < 0.001; immobile time: p < 0.001) and stressed WT mice (immobile episodes: p = 0.043, **Fig. 2E**; immobile time: p = 0.036, **Fig. 2F**). These results suggest that chronic restraint stress produces a distinct behavioral profile in DS mice on the EZM characterized by increased locomotion, head dips, and mobility, which is not observed in the stressed WT mice.

The BNST receives widespread glutamatergic and GABAergic inputs from cortical, limbic, and brainstem regions (Stamatakis et al. 2014). To assess whether chronic stress differentially alters these inputs in DS, we performed whole-cell patch-clamp recordings on stress-activated corticotropin releasing factor (CRF) neurons in the oval BNST following the 8 day restraint period (**Fig. 1A**). By utilizing the CRH reporter line, we were able to selectively patch from CRF neurons that are mostly restricted to the oval nucleus of the BNST (see Materials and Methods).

Using a Cs-methanesulfonate internal solution, we measured sEPSCs and sIPSCs voltage-clamped at -70 mV and +10 mV, respectively (**Fig. 3A**, **Fig. 3G**, see Materials and Methods). Stress elevated sEPSC amplitudes of CRF neurons of DS mice compared to both control DS (p < 0.0001) and stress WT groups (p = 0.006, **Fig. 3B**). Compared to their respective control groups, sEPSC frequency was also increased for both stress WT (p = 0.011) and stress DS groups (p < 0.001, **Fig. 3C**). sEPSC rise time was unchanged (**Fig. 3D**), and the area under the curve or the total charge transfer per event was greater in the stressed DS group compared to stressed WT (p = 0.011; **Fig. 3F**). Both stress groups had significantly faster sEPSC decay times compared to their respective controls (WT: p = 0.005; DS: p < 0.001, **Fig. 3E**).

**Figure 3:**
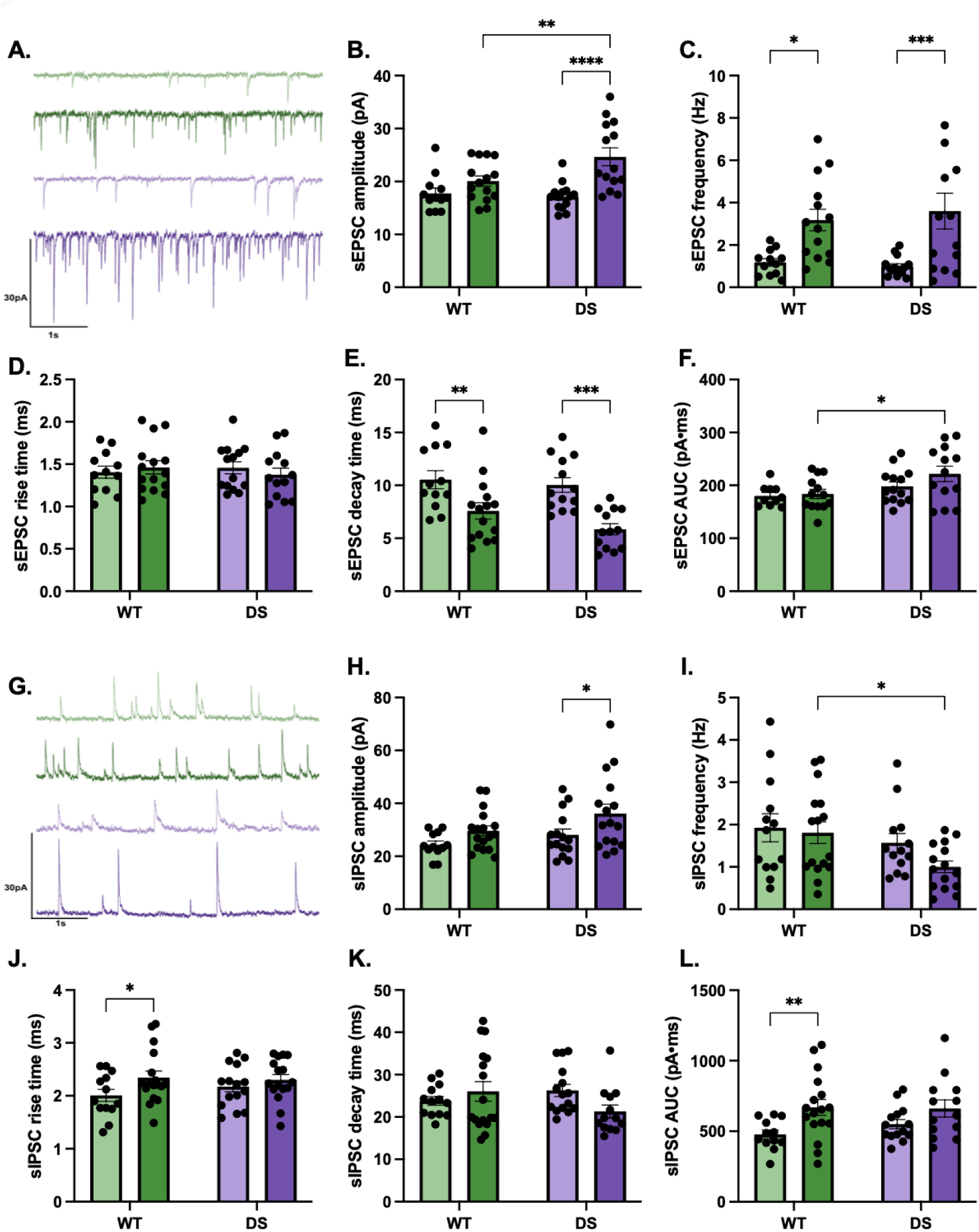
Chronic stress increases synaptic transmission onto BNST CRF neurons. **(A)** Representative traces of spontaneous excitatory postsynaptic currents (sEPSCs) recorded at −70 mV from BNST CRF neurons in WT control (light green), WT stress (dark green), DS control (light purple), and DS stress (dark purple) mice. Scale bar: 30 pA by 1 second. **(B)** sEPSC amplitude (genotype: F(1, 51) = 2.94, p = 0.092; stress: F(1, 51) = 18.02, p < 0.0001; genotype × stress interaction: F(1, 51) = 4.97, p = 0.03). **(C)** sEPSC frequency (genotype: F(1, 50) = 0.049, p = 0.83; stress: F(1, 50) = 19.29, p < 0.0001; genotype × stress interaction: F(1, 50) = 0.34, p = 0.56). **(D)** sEPSC rise time. No significant differences were observed across groups (genotype: F(1, 50) = 0.056, p = 0.81; stress: F(1, 50) = 0.026, p = 0.87; genotype × stress interaction: F(1, 50) = 0.83, p = 0.37). **(E)** sEPSC decay time (genotype: F(1, 49) = 2.40, p = 0.13; stress: F(1, 49) = 24.48, p < 0.0001; genotype × stress interaction: F(1, 49) = 0.74, p = 0.40). **(F)** sEPSC area under the curve (AUC), reflecting average charge transfer per event (genotype: F(1, 46) = 7.20, p = 0.010; stress: F(1, 46) = 1.73, p = 0.20; genotype × stress interaction: F(1, 46) = 0.92, p = 0.34). **(G)** Representative traces of spontaneous inhibitory postsynaptic currents (sIPSCs) recorded at +10 mV from BNST CRF neurons. Scale bar: 30 pA by 1 second. **(H)** sIPSC amplitude (genotype: F(1, 56) = 4.24, p = 0.044; stress: F(1, 56) = 7.12, p = 0.010; genotype × stress interaction: F(1, 56) = 0.31, p = 0.58). **(I)** sIPSC frequency (genotype: F(1, 53) = 5.67, p = 0.021; stress: F(1, 53) = 1.98, p = 0.16; genotype × stress interaction: F(1, 53) = 0.84, p = 0.36). **(J)** sIPSC rise time (genotype: F(1, 57) = 0.31, p = 0.58; stress: F(1, 57) = 4.26, p = 0.044; genotype × stress interaction: F(1, 57) = 0.88, p = 0.35). **(K)** sIPSC decay time (genotype: F(1, 54) = 0.43, p = 0.52; stress: F(1, 54) = 0.61, p = 0.44; genotype × stress interaction: F(1, 54) = 4.19, p = 0.046). **(L)** sIPSC AUC (genotype: F(1, 52) = 0.43, p = 0.52; stress: F(1, 52) = 9.27, p = 0.004; genotype × stress interaction: F(1, 52) = 0.66, p = 0.42). N = 4-6 animals per group, n = 12-17 cells per group. Statistical significance was determined by 2-way ANOVA followed by Fisher’s LSD post hoc test. *p < 0.05, **p < 0.01, ***p < 0.001, ****p < 0.0001.

Spontaneous inhibitory postsynaptic currents (sIPSC) were measured at +10 mV, and sIPSC amplitudes were increased in the stressed DS group compared to controls (p = 0.023, **Fig. 3H**), and sIPSC frequency was lower for the stressed DS group compared to stressed WT (p = 0.018, **Fig. 3I**). The inhibitory events of the stressed WT neurons exhibited faster rise times (p = 0.041, **Fig. 3J**), and a greater average charge transfer than the control WT group (p = 0.01, **Fig. 3L**). sIPSC event decay time was not affected (**Fig. 3K**).

Finally, stress differentially affected the passive membrane properties of BNST CRF neurons depending on the genotype, with a decreased membrane resistance in the stressed DS group (p = 0.041, **Table T1A**) and increased total cell capacitance in the stressed WT group (p < 0.001, **Table T1B**).

Together, these findings indicate that chronic restraint stress enhances spontaneous excitatory synaptic drive onto BNST CRF neurons in both genotypes, with DS mice showing a pronounced increase in sEPSC amplitude. Conversely, changes in inhibitory transmission appear to be more selective, with the BNST CRF neurons of stressed DS mice showing an increased sIPSC amplitude, but attenuated in other kinetics relative to WT mice.

To further investigate the altered excitatory transmission onto BNST neurons in response to stress, we measured field potential recordings and single-cell evoked synaptic properties while blocking inhibitory GABAergic transmission using picrotoxin (**Fig. 4A**, **Fig. 4D**). First, we examined whether chronic stress differentially affected overall synaptic plasticity by recording fEPSPs in the BNST. LTP was quantified as the average percent potentiation over the final 10 minutes of recording (**Fig. 4A**). Stress blunted LTP in the BNST of both WT (p = 0.007) and DS groups compared to controls (p = 0.027, **Fig. 4B**). The baseline paired pulse ratio (PPR) or release probability was not affected (**Fig. 4C**). Normalized fEPSP input-output curves were similar between all groups.

**Figure 4:**
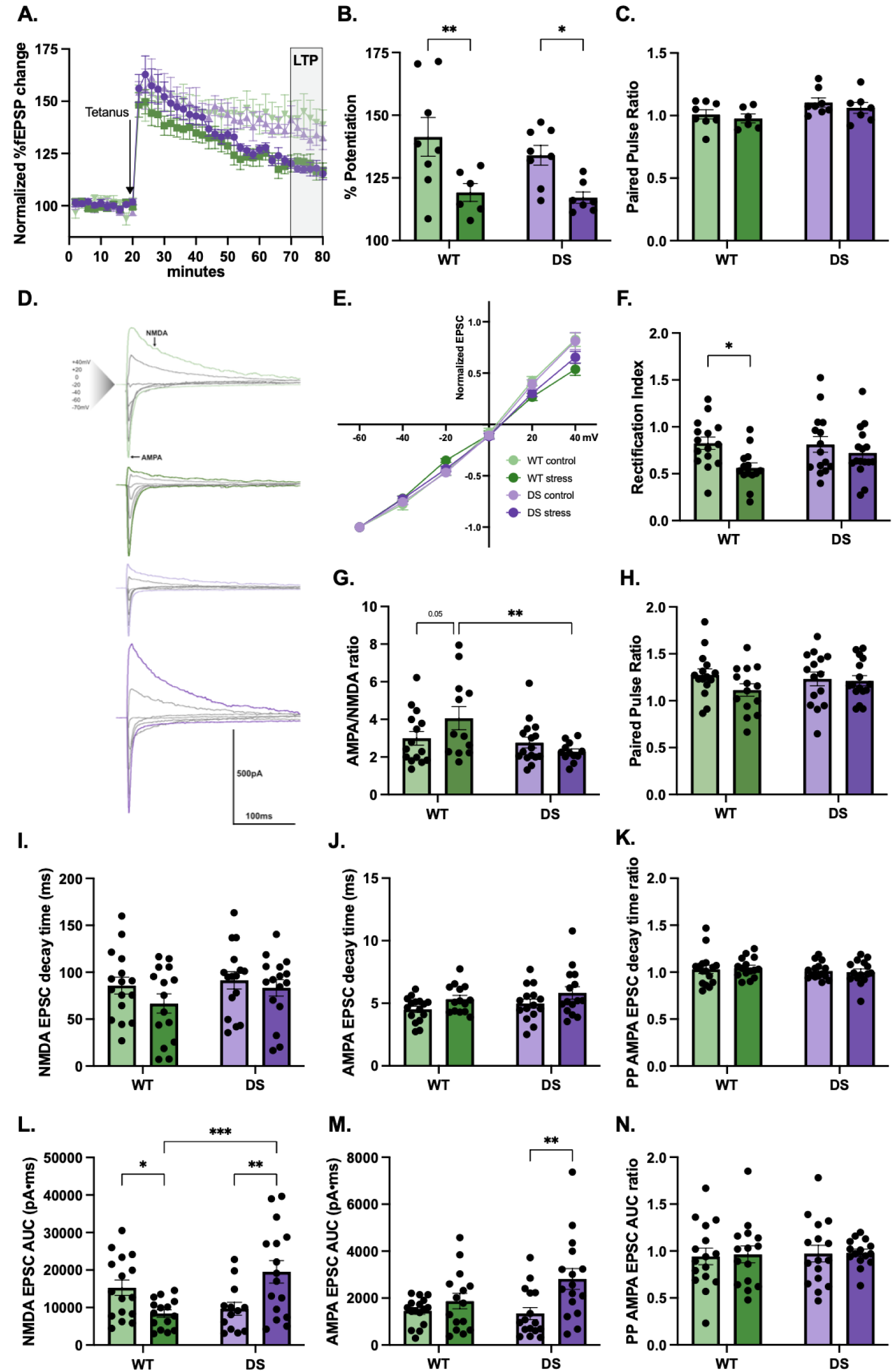
Chronic restraint stress alters synaptic plasticity and postsynaptic glutamatergic signaling in BNST CRF neurons. **(A)** Time course of normalized field excitatory postsynaptic potential (fEPSP) amplitude as a percentage change from baseline, recorded in the BNST before and after tetanic stimulation (delivered at 20 minutes, indicated by arrow) in WT control (light green), WT stress (dark green), DS control (light purple), and DS stress (dark purple) mice. The shaded region (70–80 minutes) indicates the window used to quantify long-term potentiation (LTP). **(B)** LTP quantified as percent potentiation averaged over the final 10 minutes of recording (genotype: F(1, 25) = 0.82, p = 0.37; stress: F(1, 25) = 14.13, p < 0.001; genotype × stress interaction: F(1, 25) = 0.26, p = 0.62). **(C)** Paired pulse ratio of the fEPSP sweeps (genotype: F(1, 25) = 5.36, p = 0.03; stress: F(1, 25) = 0.96, p = 0.34; genotype × stress interaction: F(1, 25) = 0.02, p = 0.89). **(A-C)** N = 6-8 animals per group. **(D)** Representative voltage-clamp traces of evoked EPSCs recorded at multiple holding potentials (−70 mV to +40 mV) from BNST CRF neurons in WT control (light green), WT stress (dark green), DS control (light purple), and DS stress (dark purple) mice, with NMDA and AMPA receptor-mediated components indicated. Scale bar: 500 pA, 100 ms. **(E)** Current-voltage relationship of normalized evoked EPSC amplitude across holding potentials for all four groups, used to assess AMPA receptor rectification. **(F)** Rectification index, calculated as the ratio of evoked EPSC amplitude at +40 mV relative to −60 mV (genotype: F(1, 56) = 1.15, p = 0.29; stress: F(1, 56) = 6.54, p = 0.013; genotype × stress interaction: F(1, 56) = 1.58, p = 0.21). **(G)** AMPA/NMDA ratio (genotype: F(1, 51) = 6.95, p = 0.011; stress: F(1, 51) = 0.58, p = 0.45; genotype × stress interaction: F(1, 51) = 4.16, p = 0.046). **(H)** Paired pulse ratio of evoked EPSC amplitude. No significant differences were observed across groups (genotype: F(1, 56) = 0.18, p = 0.67; stress: F(1, 56) = 1.95, p = 0.17; genotype × stress interaction: F(1, 56) = 1.21, p = 0.28). **(I)** NMDA receptor-mediated EPSC decay time. No significant differences were observed across groups (genotype: F(1, 59) = 1.44, p = 0.23; stress: F(1, 59) = 2.12, p = 0.15; genotype × stress interaction: F(1, 59) = 0.36, p = 0.55). **(J)** AMPA receptor-mediated EPSC decay time (genotype: F(1, 56) = 1.75, p = 0.19; stress: F(1, 56) = 5.38, p = 0.024; genotype × stress interaction: F(1, 56) = 0.002, p = 0.96). **(K)** Paired pulse ratio of AMPA EPSC decay time. No significant differences were observed across groups (genotype: F(1, 57) = 0.90, p = 0.35; stress: F(1, 57) = 0.003, p = 0.96; genotype × stress interaction: F(1, 57) = 0.17, p = 0.68). **(L)** NMDA EPSC area under the curve (AUC), reflecting total charge transfer per NMDA receptor-mediated event (genotype: F(1, 56) = 1.64, p = 0.21; stress: F(1, 56) = 0.47, p = 0.50; genotype × stress interaction: F(1, 56) = 15.20, p < 0.001). **(M)** AMPA EPSC AUC. Stress significantly increased AMPA EPSC AUC in DS mice but not WT mice (genotype: F(1, 59) = 1.76, p = 0.19; stress: F(1, 59) = 9.17, p = 0.004; genotype × stress interaction: F(1, 59) = 2.85, p = 0.10). **(N)** Paired pulse ratio of AMPA EPSC AUC. No significant differences were observed across groups (genotype: F(1, 58) = 0.09, p = 0.77; stress: F(1, 58) = 0.038, p = 0.85; genotype × stress interaction: F(1, 58) = 0.007, p = 0.93). **(F-N)** N = 4 animals per group, with n = 13-16 cells per group. Statistical significance was determined by 2-way ANOVA followed by Fisher’s LSD post hoc test. *p < 0.05, **p < 0.01, ***p < 0.001.

Next, we narrowed our focus to evoked glutamatergic transmission onto oval BNST CRF neurons to characterize postsynaptic properties. We measured AMPA receptor-mediated evoked EPSC at -70 mV and the NMDA receptor mediated evoked EPSC at +40 mV, then calculated the AMPA/NMDA ratio (**Fig. 4D**). Stress slightly increased the AMPA/NMDA ratio in the WT group compared to control WT (p = 0.055) and significantly increased compared to stressed DS (p = 0.003, **Fig. 4G**). Then, we assessed the composition of calcium-permeable AMPA receptors (CP-AMPARs) by measuring the rectification index as the ratio of evoked EPSC amplitude at +40 mV relative to -60 mV, and a lower index indicates greater inward rectification and a higher proportion of CP-AMPARs (**Fig. 4D**, **Fig. 4E**). We found decreased rectification index in only the stressed WT group compared to control WT (p = 0.01, **Fig. 4F**). The effect of stress on the average charge transferred per NMDA mediated EPSC event was bidirectional, as the AUC was decreased in the stress WT group (p = 0.024) but greater in the stress DS group (p = 0.002) compared to their respective controls (**Fig. 4L**). Only the stressed DS group showed a greater AMPA-mediated EPSC event AUC (p = 0.001, **Fig. 4M**). Both the NMDA and AMPA-mediated EPSC event decay times were unchanged across all groups (**Fig. 4I, 4J**). Similar to the fEPSP findings, the paired pulse ratio of the evoked EPSC amplitude, AMPA-mediated EPSC decay time or total charge transfer was not affected (**Fig. 4H**, **Fig. 4K**, **Fig. 4N**).

## Discussion

The impact of chronic stress on physiological, behavioral, and synaptic outcomes in Dravet syndrome remains an important but understudied topic. We demonstrated that DS mice exhibit a distinct response to chronic stress characterized by an elevated corticosterone response, increased hyperactivity and anxiety-like behavior, and synaptic changes in stress responsive neurons. These findings suggest that chronic stress may worsen physiological and behavioral outcomes in Dravet syndrome, in part through circuit changes in a brain region that is activated by seizures and stress. Our study notably focuses on the effect of stress during the critical period when spontaneous seizures first appear (Kalume et al. 2013). While prior studies have examined the impact of stress on seizures, few have investigated these effects during this specific developmental window.

### HPA Axis Dysregulation and Stress-Associated Mortality in DS Mice

Patients with epilepsy have increased cortisol reactivity to stressors which is correlated to poor seizure control, suggesting a fundamental difference in HPA axis regulation (Allendorfer et al. 2014). In agreement with these findings, we discovered that chronic restraint stress produced divergent CORT responses in WT and DS mice.

Nonstressed mice showed a reduction in CORT levels, likely reflecting heightened stress reactivity in younger mice, as the weaning process can be a stressful experience (Kikusui et al. 2006).

Stressed WT mice maintained stable CORT levels, indicating a sustained HPA axis engagement. In contrast, stressed DS mice showed a significant increase in CORT levels from baseline, suggesting an escalating stress response. Other DS mouse models show an intact acute corticosterone (CORT) response to stress (Sawyer et al. 2016). Despite this preserved acute response, the increased CORT levels observed in stressed DS mice suggest impaired HPA axis adaptation to chronic stress exposure.

This may reflect altered glucocorticoid receptor (GR) signaling expressed by inhibitory interneurons located in upstream regions that project to the BNST, along with weakened negative feedback to the HPA axis. As DS is mainly caused by a loss-of-function mutation of the Scn1a gene that encodes the alpha subunit of the Nav1.1 voltage-gated sodium channels primarily located on inhibitory interneurons, this dysfunction may be further exacerbated by excessive stress-induced activation of GRs (Yan et al. 2021; Ricobaraza et al. 2019; Kaneko et al. 2022; O’Neal et al. 2022; Perlman et al. 2021).

Restraint stress in mice with mutations in the Scn1a gene has been shown to lower seizure thresholds (Sawyer et al. 2016), suggesting that stress may increase seizure frequency and severity, subsequently increasing risk of SUDEP. Although we did not directly measure the effect of repeated stressors on seizure frequency, we found an increased mortality in stressed female DS mice. A sex-dependent difference in mortality has been reported in DS mice, potentially due to baseline differences in inhibitory interneuron tone between males and females (Fabian et al. 2024; Ellis and Honeycutt 2021; Niibori et al. 2020).

The pronounced effect of stress on epilepsy aligns with clinical observations that stress or altered mood states are frequently reported prior to SUDEP (Wood et al. 2022; Earnest et al. 1992). Additionally, epilepsy has been conceptualized as a chronic stress model and is highly impacted by stress through glucocorticoid signaling (Cano-López and González-Bono 2019; Druzhkova et al. 2022; Gulyaeva 2021). Together, this suggests a critical interaction between stress and seizures, in which stress may not only increase seizure occurrence but also the likelihood that seizures are accompanied by cardiorespiratory dysfunction.

### Stress increases hyperactivity and anxiety-like behavior in DS mice

Chronic stress produced markedly different genotype-dependent behavioral profiles. Stressed WT mice showed increased open arm time, consistent with rodent models of early life stress (Sturman et al. 2021; Toledo-Rodriguez and Sandi 2011). Similar behavioral effects have been reported following chronic corticosterone administration, which is associated with blunted LTP in the BNST (Conrad et al. 2011).

In contrast, stressed DS mice displayed increased locomotion in both open and closed arms, increased head dip frequency, and reduced immobility, suggesting hyperactivity in response to chronic stress. Rather than a typical adaptive response to repeated stress, this profile is similar to the neuropsychiatric phenotype of DS, which includes hyperactivity, impulsivity, and emotional dysregulation (Ricobaraza et al. 2019; Verbeek et al. 2015).

These maladaptive behavioral findings may reflect alterations in neurocircuitry involved in stress processing. The BNST is an extended amygdala region critically involved in stress adaptation, anxiety-like behavior, and a prime target for potential neuroadaptations (Lebow and Chen 2016; van de Poll et al. 2023; Avery et al. 2016). Given its role, these physiological findings may reflect altered BNST circuitry contributing to maladaptive responses to chronic stress. To investigate a potential circuit-level mechanism underlying these behavioral phenotypes, we examined the synaptic transmission of BNST CRF neurons.

### Alterations in spontaneous synaptic transmission to BNST CRF neurons

The BNST integrates stress and autonomic signals and sends outputs to multiple brainstem and limbic regions implicated in anxiety-related behavior and HPA axis regulation (Daniel and Rainnie 2016; L. Tran et al. 2014; S.-Y. Kim et al. 2013; Choi et al. 2007; Cheng et al. 2026). Due to the functional and structural heterogeneity of the BNST, we specifically focused on synaptic transmission onto stress-activated CRF neurons, a population that coordinates many of these processes. The BNST contains one of the highest concentrations of non-hypothalamic CRF neurons, and they receive dense excitatory input from many cortical and limbic regions, inhibitory input mainly from the central amygdala, and neuromodulatory input from brainstem and midbrain regions, placing them at a key point for investigating how stress and seizures are integrated within this region (Stamatakis et al. 2014; Luchsinger et al. 2021; Zheng et al. 2024; Chudoba and Dabrowska 2023; Wang et al. 2016). Additionally, chronic stress has been shown to change CRF expression and signaling in the BNST (Maita et al. 2022; Partridge et al. 2016; Dabrowska, Hazra, Guo, DeWitt, et al. 2013).

Notably, there is very little Scn1a expressed in the BNST, likely due to the lack of parvalbumin-expressing interneurons in the region (Xia et al. 2022). This suggests that circuit dysfunction of the BNST in DS may arise less from intrinsic deficits within the BNST and more from an altered synaptic drive from upstream inputs that are directly affected by the Scn1a mutation (Mattis et al. 2022).

We did not observe major baseline differences in spontaneous synaptic measures between the CRF neurons of nonstressed WT and DS mice. Instead, genotype-dependent differences emerged with chronic stress, indicating that stress may drive or unmask synaptic alterations. Chronic stress increased spontaneous glutamatergic event frequency and decreased the sEPSC decay time in both genotypes. The faster decay time may reflect altered AMPA receptor subunit composition (Koike-Tani et al. 2005). In the stressed WT group, unchanged sEPSC amplitude and faster decay times suggest a shift in receptor composition without substantial changes in the receptor number or synaptic localization. In contrast, the stressed DS group showed both increased sEPSC amplitude and faster decay time, and these observations may be due to various factors such as changes in receptor quantity, synaptic localization, or glutamatergic input strength. Overall, these findings suggest the genotype-dependent effects of chronic stress on postsynaptic excitatory receptors in BNST CRF neurons.

Inhibitory signaling was also altered by stress. Similar to the sEPSC amplitude change, the sIPSC amplitude was increased in the stressed DS group. The WT group did not see a change in sIPSC amplitude or event frequency, but the inhibitory events had a faster rise time and increased average charge transfer per event. This slight increase in sIPSC amplitude may reflect compensatory strengthening of postsynaptic response to inhibitory events, potentially in response to reduced inhibitory input frequency. Together, these findings suggest chronic stress remodels both excitatory and inhibitory synaptic drive onto BNST CRF neurons, with a more profound change observed in DS.

It should be noted that spontaneous recordings were obtained without synaptic blockers, whereas evoked EPSC recordings were performed in the presence of picrotoxin. Therefore, these approaches may capture distinct features of the BNST CRF circuit function.

### Alterations in evoked synaptic properties and plasticity

We observed that stress induced significant changes in excitatory transmission in BNST CRF neurons, so we further explored its effects on evoked excitatory transmission in these neurons, which is known to be differentially modulated by stress in the BNST (Conrad et al. 2011; Maita et al. 2022; Fetterly et al. 2019).

In line with findings of enhanced postsynaptic excitatory transmission, we found that stress significantly increased charge transfer of AMPA and NMDA-mediated EPSCs in the DS mice, without significant changes in kinetics. There was no change in the AMPA/NMDA ratio or rectification index in stressed DS animals. This pattern indicates proportional scaling of NMDA and AMPA receptor-mediated currents, resulting in an overall increase in total synaptic strength while preserving the relative receptor composition. These findings suggest an increase in the number of functional synapses rather than an altered receptor composition, and may reflect a failure of the DS group to undergo the adaptive synaptic remodeling observed in the WT group in response to stress.

In the stressed WT group, the decreased rectification index suggests an increase in calcium-permeable AMPA receptors (CP-AMPARs), which is supported by a trend towards a slightly elevated AMPA/NMDA ratio and a significantly reduced NMDA-mediated EPSC AUC. These findings are consistent with repeated restraint stress induced blunting of stimulation induced LTP selectively in type III BNST neurons, which include many CRF neurons (Dabrowska, Hazra, Guo, Li, et al. 2013). Our findings suggest a stress-induced strengthening of glutamatergic excitatory synapses, specifically the downregulation of GluA2-containing calcium-impermeable AMPARs.

Interestingly, the paired pulse ratio of AMPA-mediated EPSCs that showed presynaptic release probability remained unchanged, despite the increased frequency of sEPSC events. This apparent discrepancy may reflect the fact that electrically evoked EPSCs recruit a more restricted and spatially biased subset of synapses than spontaneous recordings. Given the heterogeneous distribution of local interneurons and diverse afferent inputs to BNST neurons, the evoked responses may not fully represent the broader synaptic population sampled during spontaneous event measurements.

Importantly, despite these divergent effects on basal synaptic strength, stress blunted LTP induced by high-frequency stimulation in both genotypes. This suggests that while basal excitatory transmission is regulated differentially by stress in WT and DS mice in BNST CRF neurons, the capacity for synaptic potentiation is impaired across both groups. Such stress induced LTP impairment has been previously reported in other limbic regions such as the hippocampus, and our results extend this to the BNST (E. J. Kim et al. 2015). The LTP impairment in the stress DS group is particularly notable given an elevated basal excitatory synaptic strength in this condition (Yan et al. 2021), suggesting that BNST synapses in these animals may already be operating near their ceiling potentiation, potentially limiting adaptive plasticity to future challenges and reducing flexibility under repeated stress.

At a broader circuit level, these findings raise the possibility that chronic stress may shift the functional state of BNST CRF circuitry in a maladaptive manner in Dravet syndrome. Given that DS is associated with altered excitatory to inhibitory balance in cortical and hippocampal regions, repeated seizures combined with chronic stress may increase excitatory drive onto BNST CRF neurons and promote maladaptive plasticity. Because these neurons contribute to anxiety-like behavior, stress responsivity and resilience, further investigation could provide a mechanism linking chronic stress exposure to behavioral abnormalities in DS, and potentially the neuropsychiatric comorbidities in epilepsy.

## Conclusion

Our findings reveal that chronic stress worsens outcomes in Dravet syndrome through at least two complementary mechanisms: a failure of HPA axis habituation with maladaptive behavioral phenotypes, and a circuit specific shift in the BNST suggesting impaired adaptability. As a key modulator of sustained stress responses and a region involved in seizure-induced respiratory arrest and electrophysiologically altered in DS mice (Yan et al. 2021; Xia et al. 2022), the BNST is well positioned to link chronic stress to worsened seizure and mortality outcomes in Dravet syndrome.

Several limitations of this study should be acknowledged. First, although we observed a significant sex difference in stress-induced mortality in DS mice, our electrophysiology and behavior cohorts were not powered to detect sex differences within these measures. The robust stress response in both genotypes may have masked sex differences, and future studies with larger cohorts will be needed to address this potential effect. Second, it remains unclear whether the observed synaptic changes are restricted to the CRF neurons or reflect a more generalized response in the BNST. Comparisons between CRF and non-CRF neuronal populations within the BNST under stress conditions will help refine the cellular specificity of these electrophysiological changes. More importantly, BNST CRF neurons include both local interneuron populations and projection neurons that target multiple brainstem regions (Hammack et al. 2021). Disentangling stress-induced changes across these distinct subtypes will be critical for defining circuitry changes and their potential downstream effects. Third, the young age of the mice in this study coincides with the period of interneuron maturation, which are impacted by the Scn1a+/- loss-of-function mutation in DS (Kaneko et al. 2022; Lim et al. 2018; Goff et al. 2023; Yuan et al. 2019). It is possible that the stress-induced changes we observed in this study may reflect an interaction between impaired interneuron development, stress-related neuroplasticity, and developmental neuroplasticity, rather than representing a fixed circuit.

Finally, inhibiting outflow from the dlBNST improves survival following seizure-induced respiratory arrest (Xia et al. 2022). Future studies should investigate whether chronic stress exacerbates this seizure-induced respiratory arrest in DS mice, and whether targeted modulation of BNST CRF neurons can mitigate the maladaptive effects of chronic stress in this model. Clinically, these findings underscore the importance of effective stress management in DS patients, and highlight the BNST as a promising therapeutic target in reducing the psychiatric and mortality burden in Dravet syndrome.

## Materials and methods

### Sex as a biological variable

Both male and female mice were used for this study with no discrimination, and sex differences were reported only if there was significance.

### Experimental mice

All experiments conducted with live mice were reviewed and approved by the Vanderbilt University Institutional Animal Care and Use Committee. Animal care and experimental procedures were carried out in accordance with the National Institutes of Health Guide for the Care and Use of Laboratory Animals. Mice were housed in cages containing four to five mice of the same sex and age in a mouse facility under standard laboratory conditions (14 hours light/10 hours dark cycle) and had access to food and water ad libitum.

Scn1atm1Kea mice were obtained (Miller et al. 2014). As previously described by our lab, heterozygous 129.Scn1a+/- mice were crossed with wild-type (WT) C57BL/6J mice to generate either F1 WT or heterozygous Scn1a+/- offspring used in this experiment (Yan et al. 2021). These offspring were ear tagged and genotyped between postnatal day 14 (P14) and P18, and both Dravet syndrome (DS) model (Scn1a+/-) and WT littermate mice were used in this experiment.

B6(Cg)-Crh^tm1(cre)Zjh/J^ (CRH-ires-CRE; Jackson Laboratory strain 012704) and B6.CgGt(ROSA)26Sor^tm14(CAG−TdTomato)Hze/J^ (Ai14; Jackson Laboratory strain 007914) mice were crossed to produce Crh-IRES-Cre::Ai14 offspring. These mice were subsequently crossed with 129.Scn1a+/- mice to generate F1 heterozygous Scn1a+/- (DS) or WT mice that expressed both cre recombinase and Ai14. These offspring were ear tagged and genotyped between P14 and P18, and both DS and WT littermate mice were used.

### Chronic restraint stress and body weight monitoring

Chronic restraint stress (CRS) is a well-established model to induce chronic stress in mice, and approximately one week of CRS has been shown to elicit anxiety-like behaviors (I. Tran and Gellner 2023; Prevot et al. 2025; Ye et al. 2024).

Mice aged P18 to P22 were randomly assigned to a CRS or control group, matched by sex and genotype. The CRS group underwent 8 consecutive days of restraint from 18:00 to 20:00 CST in a well-ventilated 50 mL Falcon conical tube containing two additional 1.5 mL microcentrifuge tubes to limit excessive movement during the restraint period. All mice were weighed daily throughout this 8 day period. The normalized body weight gain was calculated as the change from baseline on day 1, divided by the day 1 weight. The experimental timeline is displayed in **Figure 1A**.

### Elevated zero maze and anxiety-like behavior

The elevated zero maze (EZM) is a well established test to assess anxiety-like behavior in rodents (Braun et al. 2011). The EZM apparatus consists of a circular platform (48 cm inner diameter, 61 cm outer diameter, and 5 cm wide lanes) with two open and two closed quadrants, and elevated 50 cm off the ground. At the start of each trial, mice were placed in the center of one of the open quadrants. Between subjects, the apparatus was cleaned with 30% ethanol. An overhead camera recorded each session, and the ANY-Maze software (San Diego Instruments, San Diego, CA) was used to quantify time spent in open and closed arms, distance traveled, and number of immobile episodes.

Head dips were manually scored by two independent blinded observers and were defined as the mouse extending its body over the edge of the open arm, resulting in a distinct downward movement of the head towards the floor.

### Blood collection and serum corticosterone assay

To assess serum corticosterone (CORT) levels, mice were briefly immobilized in a 50mL conical tube and approximately 30 uL of tail blood was collected by snipping 1 mm of the distal tail. The tail snip method was selected because prior studies have shown that it produces the lowest stress-induced CORT elevation in young mice (S. Kim et al. 2018). Bleeding was stopped by applying gentle pressure, and the entire procedure was completed in under 3 minutes.

Blood was collected at 20:00 CST on the first day immediately after the first restraint or control session and on the final day of the CRS or control protocol. After allowing the blood to clot for 30 minutes, the samples were centrifuged at 1200 × g for 12 minutes. The supernatant was collected as serum and stored at -80°C. Serum CORT concentrations were quantified using the mouse/rat corticosterone ELISA kit from ALPCO (55-CORMS-E01) and levels were calculated using a 4-Parameter Logistic curve fit, according to the manufacturer’s protocol.

### Electrophysiology slice preparation

Acute brain slices were prepared from mice between P26 and P30 following the end of the 8 day CRS or control protocol. Mice were anesthetized under isoflurane and decapitated, and their brains were quickly removed and placed in an ice-cold sucrose-ACSF (in mM: 85 NaCl, 2.5 KCl, 1.25 NaH_2_PO_4_, 25 NaHCO_3_, 75 sucrose, 25 glucose, 10 DL-APV, 0.1 kynurenate, 0.5 Na L-ascorbate, 0.5 CaCl_2_, and 4 MgCl_2_). The sucrose-ACSF was continuously oxygenated with 95% O_2_/5% CO_2_. Coronal brain slices (300μm) containing the BNST were sectioned using a vibratome (Leica VT1200S, Wetzlar, Germany).

Slices were then transferred to a holding chamber containing sucrose-ACSF warmed to 37°C and slowly returned to room temperature over the course of 30 minutes. Then they were transferred to room temperature oxygenated ACSF (in mM: 125 NaCl, 2.4 KCl, 1.2 NaH_2_PO_4_, 25 NaHCO_3_, 25 glucose, 2 CaCl_2_, and 1 MgCl_2_) and maintained for at least 30 minutes before recording. This protocol has been previously utilized in our lab (Yan et al. 2021).

### Electrophysiology single cell recordings

The slices were transferred to a submerged recording chamber continuously perfused at 2.0 mL/min with oxygenated ACSF maintained at 30 ± 2°C. Neurons were identified using infrared differential interference contrast microscope (Slicescope II, Scientifica). Whole-cell patch-clamp recordings were performed using glass micropipettes with tip resistances between 4 and 6 MΩ. Signals were acquired using the Axon Multiclamp 700B amplifier (Molecular Devices), and data were sampled at 10kHz with no low-pass filter applied. Series resistance was uncompensated and data were recorded using pClamp 11 (Molecular Devices).

To record spontaneous activity, voltage-clamp was performed using a cesium-methanesulfonate based internal solution (in mM: 135 Cs-methanesulfonate, 10 KCl, 1 MgCl_2_, 20 sodium phosphocreatine, 5 QX-314, 0.2 EGTA, 4.0 Mg-ATP, and 0.3 Na-GTP (pH 7.3, 290 mOsm)). Upon breaking in, cells were held at a membrane potential of -70 mV for at least 2 minutes to confirm cell health and seal stability. Access resistance (5-32 MΩ) was monitored continuously throughout the recording, and cells showing greater than 20% change from the initial value were excluded from analysis.

Spontaneous excitatory postsynaptic currents (sEPSCs) were recorded by holding the cell at −70 mV for 5 minutes. The membrane potential was subsequently ramped to +10 mV, and spontaneous inhibitory postsynaptic currents (sIPSCs) were recorded for 5 minutes. Analysis was performed on 120 seconds of stable recording, and events were detected using Easy Electrophysiology.

To isolate evoked EPSCs, picrotoxin (dissolved in DMSO) was added to the ACSF to achieve a final concentration of 100 μM, to block both synaptic and extrasynaptic GABAA receptors. Evoked glutamatergic EPSCs were recorded by electrical stimulations delivered by a monopolar glass electrode placed dorsal to the recorded neuron and controlled by a SIU91 constant current isolator (Cygnus Tech). Input-output curves were generated by stimulating the slice at intensities of 10, 25, 50, 75, and 100 uA. The stimulation intensity that produced 50% of the maximum EPSC amplitude was used for all subsequent recordings.

AMPA/NMDA ratios were calculated based on the peak AMPAR-mediated EPSC amplitude measured at −70 mV and the NMDAR-mediated EPSC component measured at 40 ms after electrical stimulation at +40 mV. The paired-pulse ratio (PPR) was determined as the ratio of EPSC2/EPSC1 at a 40 ms interstimulus interval while the cell was voltage clamped at -70 mV. The rectification was assessed by measuring peak EPSC amplitudes at holding potentials ranging from -60 mV to +40 mV in 20 mV increments. The rectification index was calculated as the ratio of the peak amplitudes at +40 mV/-60 mV.

### Electrophysiology field potential recordings

Slices were perfused with oxygenated ACSF containing 100 uM picrotoxin in a 32°C recording chamber. Field EPSPs (fEPSP) were evoked using a bipolar stainless steel stimulating electrode, and a borosilicate glass recording electrode filled with ACSF was placed ventral to the stimulating site.

Input-output curves were first generated to determine the stimulation intensity that elicited approximately 40% of the maximum fEPSP response. A stable baseline was then recorded for at least 20 minutes. Long term potentiation (LTP) was induced using two trains of 100Hz, 1 second tetanus at 20 second intertrain intervals. LTP was quantified as the average N2 amplitude 50 to 60 minutes post-tetanus, normalized to baseline. Recordings were excluded if the N1 amplitude changed by more than 20%.

## Statistical analysis

All electrophysiology data were analyzed after applying a 2 kHz low-pass filter, and events were detected using either pClamp11 (Molecular Devices) or Easy Electrophysiology. All statistical analyses were performed and graphed using GraphPad Prism 11. Unless otherwise stated, all data were analyzed using repeated measures two way ANOVA followed by Fisher’s LSD or false discovery rate (FDR) post hoc tests, which were conducted only when a significant main effect or interaction was observed. Survival data were analyzed using the Kaplan-Meier survival curve with a log-rank (Mantel-Cox) test. Outliers were identified and removed using the Robust regression and outlier (ROUT) method in GraphPad Prism with a Q value of 5%. Graphs show the mean ± SEM. Comprehensive statistics are presented in **Table 1**.

## Authorship contribution statement

EH and WPN conceptualized the study. EH, EYX, JGM, JQ, SMM, YV, GRB, AAM contributed to data acquisition. EH, EYX, JQ, SMM, BMM analyzed data. EH compiled all figures. EH drafted the manuscript. EH, EYX, JGM, LD, JAM, BMM, FCV, WPN reviewed and edited the manuscript.

## Acknowledgements

Research reported in this publication was supported by the National Institute of Neurological Disorders and Stroke (NINDS) of the National Institutes of Health (NIH), under Award Number R01NS133169-03, a Research Grant from the Dravet Syndrome Foundation, and the Vanderbilt Kennedy Center Nicholas Hobbs Discovery Award. The elevated zero maze was performed in part through the use of the Mouse Neurobehavior Core lab at Vanderbilt University. The Vanderbilt Neurobehavioral Core facility receives funding from the Vanderbilt Kennedy Center P50HD103537-05.

## References

Allendorfer, Jane B., Heidi Heyse, Lucy Mendoza, et al. 2014. “Physiologic and Cortical Response to Acute Psychosocial Stress in Left Temporal Lobe Epilepsy - a Pilot Cross-Sectional fMRI Study.” Epilepsy & Behavior: E&B 36 (July): 115–23. 10.1016/j.yebeh.2014.05.003.

Avery, S. N., J. A. Clauss, and J. U. Blackford. 2016. “The Human BNST: Functional Role in Anxiety and Addiction.” Neuropsychopharmacology 41 (1): 126–41. 10.1038/npp.2015.185.

Braun, Amanda A., Matthew R. Skelton, Charles V. Vorhees, and Michael T. Williams. 2011. “Comparison of the Elevated plus and Elevated Zero Mazes in Treated and Untreated Male Sprague-Dawley Rats: Effects of Anxiolytic and Anxiogenic Agents.” Pharmacology, Biochemistry, and Behavior 97 (3): 406–15. 10.1016/j.pbb.2010.09.013.

Cano-López, Irene, and Esperanza González-Bono. 2019. “Cortisol Levels and Seizures in Adults with Epilepsy: A Systematic Review.” Neuroscience & Biobehavioral Reviews 103 (August): 216–29. 10.1016/j.neubiorev.2019.05.023.

Catalán-Aguilar, Judit, Esperanza González-Bono, and Irene Cano-López. 2025. “Perceived Stress in Adults with Epilepsy: A Systematic Review.” Neuroscience & Biobehavioral Reviews 170 (March): 106065. 10.1016/j.neubiorev.2025.106065.

Chen, Zhibin, Martin J. Brodie, Ding Ding, and Patrick Kwan. 2023. “Editorial: Epidemiology of Epilepsy and Seizures.” Frontiers in Epidemiology 3 (August): 1273163. 10.3389/fepid.2023.1273163.

Cheng, Xin, Yubo Hu, Yan Zhao, et al. 2026. “Generalization of Negative Memories Drives the Development of Psychological Distress-Related Behaviors.” The Innovation 0 (0). 10.1016/j.xinn.2026.101306.

Choi, Dennis C., Amy R. Furay, Nathan K. Evanson, Michelle M. Ostrander, Yvonne M. Ulrich-Lai, and James P. Herman. 2007. “Bed Nucleus of the Stria Terminalis Subregions Differentially Regulate Hypothalamic–Pituitary–Adrenal Axis Activity: Implications for the Integration of Limbic Inputs.” The Journal of Neuroscience 27 (8): 2025–34. 10.1523/JNEUROSCI.4301-06.2007.

Chudoba, Rachel, and Joanna Dabrowska. 2023. “Distinct Populations of Corticotropin-Releasing Factor (CRF) Neurons Mediate Divergent yet Complementary Defensive Behaviors in Response to a Threat.” Neuropharmacology 228 (May): 109461. 10.1016/j.neuropharm.2023.109461.

Conrad, Kelly L., Katherine M. Louderback, Caitlin P. Gessner, and Danny G. Winder. 2011. “Stress-Induced Alterations in Anxiety-like Behavior and Adaptations in Plasticity in the Bed Nucleus of the Stria Terminalis.” Physiology & Behavior 104 (2): 248–56. 10.1016/j.physbeh.2011.03.001.

Dabrowska, Joanna, Rimi Hazra, Ji-Dong Guo, ChenChen Li, et al. 2013. “Striatal-Enriched Protein Tyrosine Phosphatase—STEPs Toward Understanding Chronic Stress-Induced Activation of Corticotrophin Releasing Factor Neurons in the Rat Bed Nucleus of the Stria Terminalis.” *Biological Psychiatry*, Stress: Impact on Brain and Body, vol. 74 (11): 817–26. 10.1016/j.biopsych.2013.07.032.

Dabrowska, Joanna, Rimi Hazra, Ji-Dong Guo, Sarah DeWitt, and Donald G. Rainnie. 2013. “Central CRF Neurons Are Not Created Equal: Phenotypic Differences in CRF-Containing Neurons of the Rat Paraventricular Hypothalamus and the Bed Nucleus of the Stria Terminalis.” Frontiers in Neuroscience 7 (August): 156. 10.3389/fnins.2013.00156.

Dabrowska, Joanna, Daisy Martinon, Mahsa Moaddab, and Donald G. Rainnie. 2016. “Targeting Corticotropin-Releasing Factor (CRF) Projections from the Oval Nucleus of the BNST Using Cell-Type Specific Neuronal Tracing Studies in Mouse and Rat Brain.” Journal of Neuroendocrinology 28 (12): 10.1111/jne.12442. 10.1111/jne.12442.

Daniel, Sarah E., and Donald G. Rainnie. 2016. “Stress Modulation of Opposing Circuits in the Bed Nucleus of the Stria Terminalis.” Neuropsychopharmacology 41 (1): 103–25. 10.1038/npp.2015.178.

Dhaher, Roni, Richard A. Bronen, Linda Spencer, et al. 2022. “Dorsal Bed Nucleus of Stria Terminalis in Depressed and Nondepressed Temporal Lobe Epilepsy Patients.” Epilepsia 63 (10): 2561–70. 10.1111/epi.17377.

Druzhkova, Tatyana A., Alexander A. Yakovlev, Flora K. Rider, Mikhail S. Zinchuk, Alla B. Guekht, and Natalia V. Gulyaeva. 2022. “Elevated Serum Cortisol Levels in Patients with Focal Epilepsy, Depression, and Comorbid Epilepsy and Depression.” International Journal of Molecular Sciences 23 (18): 10414. 10.3390/ijms231810414.

Earnest, M. P., G. E. Thomas, R. A. Eden, and K. F. Hossack. 1992. “The Sudden Unexplained Death Syndrome in Epilepsy: Demographic, Clinical, and Postmortem Features.” Epilepsia 33 (2): 310–16. 10.1111/j.1528-1157.1992.tb02321.x.

Ellis, Seneca N., and Jennifer A. Honeycutt. 2021. “Sex Differences in Affective Dysfunction and Alterations in Parvalbumin in Rodent Models of Early Life Adversity.” Frontiers in Behavioral Neuroscience 15 (November): 741454. 10.3389/fnbeh.2021.741454.

England, Mary Jane, Catharyn T. Liverman, Andrea M. Schultz, and Larisa M. Strawbridge. 2012a. “A Summary of the Institute of Medicine Report.” Epilepsy & Behavior : E&B 25 (2): 266–76. 10.1016/j.yebeh.2012.06.016.

England, Mary Jane, Catharyn T. Liverman, Andrea M. Schultz, and Larisa M. Strawbridge. 2012b. “Summary.” Epilepsy Currents 12 (6): 245–53. 10.5698/1535-7511-12.6.245.

Espinosa-Garcia, Claudia, Helena Zeleke, and Asheebo Rojas. 2021. “Impact of Stress on Epilepsy: Focus on Neuroinflammation—A Mini Review.” International Journal of Molecular Sciences 22 (8): 4061. 10.3390/ijms22084061.

Fabian, Carly B., Nilah D. Jordan, Rebecca H. Cole, et al. 2024. “Parvalbumin Interneuron mGlu5 Receptors Govern Sex Differences in Prefrontal Cortex Physiology and Binge Drinking.” Neuropsychopharmacology 49 (12): 1861–71. 10.1038/s41386-024-01889-0.

Fetterly, Tracy L., Aakash Basu, Brett P. Nabit, et al. 2019. “α2A-Adrenergic Receptor Activation Decreases Parabrachial Nucleus Excitatory Drive onto BNST CRF Neurons and Reduces Their Activity In Vivo.” The Journal of Neuroscience 39 (3): 472–84. 10.1523/JNEUROSCI.1035-18.2018.

Gandy, Milena, Avani C. Modi, Janelle L. Wagner, et al. 2021. “Managing Depression and Anxiety in People with Epilepsy: A Survey of Epilepsy Health Professionals by the ILAE Psychology Task Force.” Epilepsia Open 6 (1): 127–39. 10.1002/epi4.12455.

Goff, Kevin M., Sophie R. Liebergall, Evan Jiang, Ala Somarowthu, and Ethan M. Goldberg. 2023. “VIP Interneuron Impairment Promotes in Vivo Circuit Dysfunction and Autism-Related Behaviors in Dravet Syndrome.” Cell Reports 42 (6): 112628. 10.1016/j.celrep.2023.112628.

Gulyaeva, Natalia V. 2021. “Stress-Associated Molecular and Cellular Hippocampal Mechanisms Common for Epilepsy and Comorbid Depressive Disorders.” Biochemistry (Moscow) 86 (6): 641–56. 10.1134/S0006297921060031.

Gurgu, Raluca Simona, Adela Magdalena Ciobanu, Roxana Ionela Danasel, and Cristina Aura Panea. 2021. “Psychiatric Comorbidities in Adult Patients with Epilepsy (A Systematic Review).” Experimental and Therapeutic Medicine 22 (2): 909. 10.3892/etm.2021.10341.

Hammack, Sayamwong E., Karen M. Braas, and Victor May. 2021. “Chemoarchitecture of the BNST: Neurophenotypic Diversity and Function.” Handbook of Clinical Neurology 179: 385–402. 10.1016/B978-0-12-819975-6.00025-X.

Kalume, Franck, Ruth E. Westenbroek, Christine S. Cheah, et al. 2013. “Sudden Unexpected Death in a Mouse Model of Dravet Syndrome.” The Journal of Clinical Investigation 123 (4): 1798–808. 10.1172/JCI66220.

Kaneko, Keisuke, Christopher B. Currin, Kevin M. Goff, et al. 2022. “Developmentally Regulated Impairment of Parvalbumin Interneuron Synaptic Transmission in an Experimental Model of Dravet Syndrome.” Cell Reports 38 (13): 110580. 10.1016/j.celrep.2022.110580.

Kikusui, Takefumi, Kayo Nakamura, Yoshie Kakuma, and Yuji Mori. 2006. “Early Weaning Augments Neuroendocrine Stress Responses in Mice.” Behavioural Brain Research 175 (1): 96–103. 10.1016/j.bbr.2006.08.007.

Kim, Eun Joo, Blake Pellman, and Jeansok J. Kim. 2015. “Stress Effects on the Hippocampus: A Critical Review.” Learning & Memory 22 (9): 411–16. 10.1101/lm.037291.114.

Kim, Sarah, Daphne Foong, Mark S. Cooper, Markus J. Seibel, and Hong Zhou. 2018. “Comparison of Blood Sampling Methods for Plasma Corticosterone Measurements in Mice Associated with Minimal Stress-Related Artefacts.” Steroids 135 (July): 69–72. 10.1016/j.steroids.2018.03.004.

Kim, Sung-Yon, Avishek Adhikari, Soo Yeun Lee, et al. 2013. “Diverging Neural Pathways Assemble a Behavioural State from Separable Features in Anxiety.” Nature 496 (7444): 219–23. 10.1038/nature12018.

Koike-Tani, Maki, Naoto Saitoh, and Tomoyuki Takahashi. 2005. “Mechanisms Underlying Developmental Speeding in AMPA-EPSC Decay Time at the Calyx of Held.” The Journal of Neuroscience 25 (1): 199–207. 10.1523/JNEUROSCI.3861-04.2005.

Lang, Johannes D., David C. Taylor, and Burkhard S. Kasper. 2018. “Stress, Seizures, and Epilepsy: Patient Narratives.” Epilepsy & Behavior 80 (March): 163–72. 10.1016/j.yebeh.2018.01.005.

Lathers, Claire M., and Paul L. Schraeder. 2006. “Stress and Sudden Death.” Epilepsy & Behavior 9 (2): 236–42. 10.1016/j.yebeh.2006.06.001.

Lebow, M. A., and A. Chen. 2016. “Overshadowed by the Amygdala: The Bed Nucleus of the Stria Terminalis Emerges as Key to Psychiatric Disorders.” Molecular Psychiatry 21 (4): 450–63. 10.1038/mp.2016.1.

Lim, Lynette, Da Mi, Alfredo Llorca, and Oscar Marín. 2018. “Development and Functional Diversification of Cortical Interneurons.” Neuron 100 (2): 294–313. 10.1016/j.neuron.2018.10.009.

Luchsinger, Joseph R., Tracy L. Fetterly, Kellie M. Williford, et al. 2021. “Delineation of an Insula-BNST Circuit Engaged by Struggling Behavior That Regulates Avoidance in Mice.” Nature Communications 12 (1): 3561. 10.1038/s41467-021-23674-z.

Maita, Isabella, Troy A. Roepke, and Benjamin A. Samuels. 2022. “Chronic Stress-Induced Synaptic Changes to Corticotropin-Releasing Factor-Signaling in the Bed Nucleus of the Stria Terminalis.” Frontiers in Behavioral Neuroscience 16 (August): 903782. 10.3389/fnbeh.2022.903782.

Mattis, Joanna, Ala Somarowthu, Kevin M. Goff, et al. 2022. “Corticohippocampal Circuit Dysfunction in a Mouse Model of Dravet Syndrome.” eLife 11 (February): e69293. 10.7554/eLife.69293.

McKee, Heather R., and Michael D. Privitera. 2017. “Stress as a Seizure Precipitant: Identification, Associated Factors, and Treatment Options.” Seizure, 25th Anniversary Issue, vol. 44 (January): 21–26. 10.1016/j.seizure.2016.12.009.

Miller, A. R., N. A. Hawkins, C. E. McCollom, and J. A. Kearney. 2014. “Mapping Genetic Modifiers of Survival in a Mouse Model of Dravet Syndrome.” Genes, Brain, and Behavior 13 (2): 163–72. 10.1111/gbb.12099.

Mula, Marco, Andres M. Kanner, Nathalie Jetté, and Josemir W. Sander. 2021. “Psychiatric Comorbidities in People With Epilepsy.” Neurology: Clinical Practice 11 (2): e112–20. 10.1212/CPJ.0000000000000874.

Niibori, Yosuke, Shiron J. Lee, Berge A. Minassian, and David R. Hampson. 2020. “Sexually Divergent Mortality and Partial Phenotypic Rescue After Gene Therapy in a Mouse Model of Dravet Syndrome.” Human Gene Therapy 31 (5–6): 339–51. 10.1089/hum.2019.225.

O’Neal, Teri B., Sanjay Shrestha, Harsimar Singh, et al. 2022. “Sudden Unexpected Death in Epilepsy.” Neurology International 14 (3): 600–613. 10.3390/neurolint14030048.

Partridge, John G., Patrick A. Forcelli, Ruixi Luo, et al. 2016. “Stress Increases GABAergic Neurotransmission in CRF Neurons of the Central Amygdala and Bed Nucleus Stria Terminalis.” Neuropharmacology 107 (August): 239–50. 10.1016/j.neuropharm.2016.03.029.

Perlman, George, Arnaud Tanti, and Naguib Mechawar. 2021. “Parvalbumin Interneuron Alterations in Stress-Related Mood Disorders: A Systematic Review.” Neurobiology of Stress 15 (August): 100380. 10.1016/j.ynstr.2021.100380.

Poll, Yana van de, Yasmin Cras, and Tommas J. Ellender. 2023. “The Neurophysiological Basis of Stress and Anxiety - Comparing Neuronal Diversity in the Bed Nucleus of the Stria Terminalis (BNST) across Species.” Frontiers in Cellular Neuroscience 17 (August). 10.3389/fncel.2023.1225758.

Prentice, Anna J., Ian McSalley, Jan H. Magielski, et al. 2026. “Characterizing SCN1A-Related Disorders Using Real-World Data Across 681 Patient-Years.” *medRxiv*, March 2, 2026.02.24.26346493. 10.64898/2026.02.24.26346493.

Prevot, Thomas D., Jaime K. Knoch, Dipashree Chatterjee, et al. 2025. “Dynamic Behavioral and Molecular Changes Induced by Chronic Restraint Stress Exposure in Mice.” International Journal of Molecular Sciences 27 (1): 167. 10.3390/ijms27010167.

Reda, Anas, Derek J. Doss, Ghassan S. Makhoul, et al. 2025. “Altered Interictal Bed Nucleus of Stria Terminalis Connectivity in Patients With Temporal Lobe Epilepsy.” Neurology 105 (11): e214385. 10.1212/WNL.0000000000214385.

Ricobaraza, Ana, Lucia Mora-Jimenez, Elena Puerta, et al. 2019. “Epilepsy and Neuropsychiatric Comorbidities in Mice Carrying a Recurrent Dravet Syndrome SCN1A Missense Mutation.” Scientific Reports 9 (October): 14172. 10.1038/s41598-019-50627-w.

Sawyer, N. T., A. W. Helvig, C. D. Makinson, M. J. Decker, G. N. Neigh, and A. Escayg. 2016. “Scn1a Dysfunction Alters Behavior but Not the Effect of Stress on Seizure Response.” *Genes*, Brain, and Behavior 15 (3): 335–47. 10.1111/gbb.12281.

Stamatakis, Alice M., Dennis R. Sparta, Joshua H. Jennings, Zoe A. McElligott, Heather Decot, and Garret D. Stuber. 2014. “Amygdala and Bed Nucleus of the Stria Terminalis Circuitry: Implications for Addiction-Related Behaviors.” Neuropharmacology 76 (0 0): 10.1016/j.neuropharm.2013.05.046. 10.1016/j.neuropharm.2013.05.046.

Sturman, Oliver, Lukas von Ziegler, Mattia Privitera, et al. 2021. “Chronic Adolescent Stress Increases Exploratory Behavior but Does Not Appear to Change the Acute Stress Response in Adult Male C57BL/6 Mice.” Neurobiology of Stress 15 (September): 100388. 10.1016/j.ynstr.2021.100388.

Toledo-Rodriguez, Maria, and Carmen Sandi. 2011. “Stress during Adolescence Increases Novelty Seeking and Risk-Taking Behavior in Male and Female Rats.” Frontiers in Behavioral Neuroscience 5 (April): 17. 10.3389/fnbeh.2011.00017.

Tran, Inès, and Anne-Kathrin Gellner. 2023. “Long-Term Effects of Chronic Stress Models in Adult Mice.” Journal of Neural Transmission 130 (9): 1133–51. 10.1007/s00702-023-02598-6.

Tran, Lee, Jay Schulkin, and Beverley Greenwood-Van Meerveld. 2014. “Importance of CRF Receptor-Mediated Mechanisms of the Bed Nucleus of the Stria Terminalis in the Processing of Anxiety and Pain.” Neuropsychopharmacology 39 (11): 2633–45. 10.1038/npp.2014.117.

Valassina, Nicholas, Simone Brusco, Alessia Salamone, et al. 2022. “Scn1a Gene Reactivation after Symptom Onset Rescues Pathological Phenotypes in a Mouse Model of Dravet Syndrome.” Nature Communications 13 (1): 161. 10.1038/s41467-021-27837-w.

Verbeek, Nienke E., Merel Wassenaar, Jolien S. van Campen, et al. 2015. “Seizure Precipitants in Dravet Syndrome: What Events and Activities Are Specifically Provocative Compared with Other Epilepsies?” Epilepsy & Behavior: E&B 47 (June): 39–44. 10.1016/j.yebeh.2015.05.008.

Wang, Li, Minjie Shen, Changyou Jiang, Lan Ma, and Feifei Wang. 2016. “Parvalbumin Interneurons of Central Amygdala Regulate the Negative Affective States and the Expression of Corticotrophin-Releasing Hormone During Morphine Withdrawal.” International Journal of Neuropsychopharmacology 19 (11): pyw060. 10.1093/ijnp/pyw060.

Wood, Ian B., Rion Brattig Correia, Wendy R. Miller, and Luis M. Rocha. 2022. “Small Cohort of Patients with Epilepsy Showed Increased Activity on Facebook before Sudden Unexpected Death.” Epilepsy & Behavior : E&B 128 (March): 108580. 10.1016/j.yebeh.2022.108580.

Xia, Maya, Benjamin Owen, Jeremy Chiang, et al. 2022. “Disruption of Synaptic Transmission in the Bed Nucleus of the Stria Terminalis Reduces Seizure-Induced Death in DBA/1 Mice and Alters Brainstem E/I Balance.” ASN NEURO 14 (May): 17590914221103188. 10.1177/17590914221103188.

Yan, Wen Wei, Maya Xia, Jeremy Chiang, et al. 2021. “Enhanced Synaptic Transmission in the Extended Amygdala and Altered Excitability in an Extended Amygdala to Brainstem Circuit in a Dravet Syndrome Mouse Model.” eNeuro 8 (3): ENEURO.0306-20.2021. 10.1523/ENEURO.0306-20.2021.

Ye, Fan, Meng-Chen Dong, Chen-Xi Xu, et al. 2024. “Effects of Different Chronic Restraint Stress Periods on Anxiety- and Depression-like Behaviors and Tryptophan-Kynurenine Metabolism along the Brain-Gut Axis in C57BL/6N Mice.” European Journal of Pharmacology 965 (February): 176301. 10.1016/j.ejphar.2023.176301.

Yuan, Yukun, Heather A. O’Malley, Melissa A. Smaldino, Alexandra A. Bouza, Jacob M. Hull, and Lori L. Isom. 2019. “Delayed Maturation of GABAergic Signaling in the Scn1a and Scn1b Mouse Models of Dravet Syndrome.” Scientific Reports 9 (1): 6210. 10.1038/s41598-019-42191-0.

Zheng, Xianli, Li Dingpeng, Xingke Yan, Xiaoqiang Yao, and Yongrui Wang. 2024. “The Role and Mechanism of 5-HTDRN-BNST Neural Circuit in Anxiety and Fear Lesions.” Frontiers in Neuroscience 18 (May): 1362899. 10.3389/fnins.2024.1362899.

