## Supplementary material for "Chronic Stress Alters Dorsal Bed Nucleus of Stria Terminalis Synaptic Neurotransmission in a Dravet Syndrome Mouse Model": Table

| Figure | Experiment | Animal (N) and/or cell number (n). Values are shown after removing outliers with ROUT. | Statistics performed | Results |
|---|---|---|---|---|
| 1B | WT control vs WT stress body weight gain | WT control: N = 21<br>WT stress: N = 24 | Two-way repeated measures ANOVA | Time: F(1.31, 56.33) = 241.7, p < 0.0001 |
| | | | | Stress: F(1, 43) = 55.76; p < 0.0001 |
| | | | | Interaction: F(1.31, 56.33) = 48.29; p < 0.0001 |
| | | | Post hoc Benjamini–Hochberg FDR correction | Day 2: p < 0.0001 |
| | | | | Day 3: p < 0.0001 |
| | | | | Day 4: p < 0.0001 |
| | | | | Day 5: p < 0.0001 |
| | | | | Day 6: p < 0.0001 |
| | | | | Day 7: p < 0.0001 |
| | | | | Day 8: p < 0.0001 |
| 1C | DS control vs DS stress body weight gain | DS control: N = 21<br>DS stress: N = 22 | Two-way repeated measures ANOVA | Time: F(1.35, 55.32) = 121.9, p < 0.0001 |
| | | | | Stress: F(1, 41) = 29.45, p < 0.0001 |
| | | | | Interaction: F(1.35, 55.32) = 11.98, p < 0.001 |
| | | | Post hoc Benjamini–Hochberg FDR correction | Day 2: p = 0.0002 |
| | | | | Day 3: p < 0.0001 |
| | | | | Day 4: p < 0.0001 |
| | | | | Day 5: p < 0.0001 |
| | | | | Day 6: p < 0.0001 |
| | | | | Day 7: p < 0.0001 |
| | | | | Day 8: p < 0.0001 |
| 1D | WT control vs WT stress CORT | WT control: N = 10<br>WT stress: N = 10 | Two-way repeated measures ANOVA | Time: F(1, 18) = 3.94, p = 0.063 |
| | | | | Stress: F(1, 18) = 12.99, p = 0.002 |
| | | | | Interaction: F(1, 18) = 7.55, p = 0.013 |
| | | | Fisher's LSD post hoc test | Baseline control vs stress: p = 0.39 |
| | | | | Final control vs stress: p < 0.0001 |
| | | | | Control baseline vs final: p = 0.004 |
| | | | | Stress baseline vs final: p = 0.60 |
| 1E | DS control vs DS stress CORT | DS control: N = 10<br>DS stress: N = 9 | Two-way repeated measures ANOVA | Time: F(1, 17) = 0.057, p = 0.81 |
| | | | | Stress: F(1, 17) = 20.11, p < 0.001 |
| | | | | Interaction: F(1, 17) = 12.97, p = 0.002 |
| | | | Fisher's LSD post hoc test | Baseline control vs stress: p = 0.19 |
| | | | | Final control vs stress: p < 0.0001 |
| | | | | Control baseline vs final: p = 0.013 |
| | | | | Stress baseline vs final: p = 0.033 |

| | | | | |
|---|---|---|---|---|
| 1F | DS stress male vs female survival | Male DS stress: N = 19<br>Female DS stress: N = 25 | Logrank Mantel-Cox test | $\chi^2(1) = 4.45$, $p = 0.035$ |
| 2B | EZM distance travelled | WT control: N = 13<br>WT stress: N = 12<br>DS control: N = 9<br>DS stress: N = 9 | Two-way ANOVA | Interaction: $F(1, 39) = 7.70$, $p = 0.008$ |
| | | | | Genotype: $F(1, 39) = 0.53$, $p = 0.47$ |
| | | | | Stress: $F(1, 39) = 10.53$, $p = 0.002$ |
| | | | Fisher's LSD post hoc test | WT control vs stress: $p = 0.72$ |
| | | | | DS control vs stress: $p < 0.001$ |
| | | | | Control WT vs DS: $p = 0.15$ |
| | | | | Stress WT vs DS: $p = 0.019$ |
| 2C | EZM open arm time | WT control: N = 13<br>WT stress: N = 12<br>DS control: N = 9<br>DS stress: N = 9 | Two-way ANOVA | Interaction: $F(1, 39) = 3.11$, $p = 0.086$ |
| | | | | Genotype: $F(1, 39) = 0.26$, $p = 0.61$ |
| | | | | Stress: $F(1, 39) = 4.80$, $p = 0.035$ |
| | | | Fisher's LSD post hoc test | WT control vs stress: $p = 0.004$ |
| | | | | DS control vs stress: $p = 0.78$ |
| 2D | EZM head dips | WT control: N = 13<br>WT stress: N = 12<br>DS control: N = 8<br>DS stress: N = 9 | Two-way ANOVA | Interaction: $F(1, 38) = 2.81$, $p = 0.10$ |
| | | | | Genotype: $F(1, 38) = 0.93$, $p = 0.34$ |
| | | | | Stress: $F(1, 38) = 10.54$, $p = 0.002$ |
| | | | Fisher's LSD post hoc test | WT control vs stress: $p = 0.22$ |
| | | | | DS control vs stress: $p = 0.003$ |
| 2E | EZM Immobile episodes | WT control: N = 13<br>WT stress: N = 12<br>DS control: N = 8<br>DS stress: N = 9 | Two-way ANOVA | Interaction: $F(1, 38) = 4.25$, $p = 0.046$ |
| | | | | Genotype: $F(1, 38) = 0.75$, $p = 0.39$ |
| | | | | Stress: $F(1, 38) = 15.18$, $p < 0.001$ |
| | | | Fisher's LSD post hoc test | WT control vs stress: $p = 0.15$ |
| | | | | DS control vs stress: $p < 0.001$ |
| | | | | Control WT vs DS: $p = 0.41$ |
| | | | | Stress WT vs DS: $p = 0.043$ |
| 2F | EZM Immobile time | WT control: N = 13<br>WT stress: N = 12<br>DS control: N = 9<br>DS stress: N = 9 | Two-way ANOVA | Interaction: $F(1, 39) = 6.31$, $p = 0.016$ |
| | | | | Genotype: $F(1, 39) = 0.34$, $p = 0.56$ |
| | | | | Stress: $F(1, 39) = 12.40$, $p = 0.001$ |
| | | | Fisher's LSD post hoc test | WT control vs stress: $p = 0.44$ |
| | | | | DS control vs stress: $p < 0.001$ |
| | | | | Control WT vs DS: $p = 0.18$ |
| | | | | Stress WT vs DS: $p = 0.036$ |
| 3B | sEPSC amplitude | WT control: N = 4 animals, n = 12 cells<br>WT stress: N = 6 animals, n = 15 cells<br>DS control: N = 4 animals, n = 14 cells<br>DS stress: N = 4 animals, n = 14 cells | Two-way ANOVA | Interaction: $F(1, 51) = 4.97$, $p = 0.03$ |
| | | | | Genotype: $F(1, 51) = 2.94$, $p = 0.092$ |
| | | | | Stress: $F(1, 51) = 18.02$, $p < 0.0001$ |
| | | | Fisher's LSD post hoc test | WT control vs stress: $p = 0.17$ |
| | | | | DS control vs stress: $p < 0.0001$ |

| | | | | |
|---|---|---|---|---|
| | | | | Control WT vs DS: p = 0.73 |
| | | | | Stress WT vs DS: p = 0.006 |
| 3C | sEPSC frequency | WT control: N = 4 animals, n = 12 cells<br>WT stress: N = 6 animals, n = 14 cells<br>DS control: N = 4 animals, n = 14 cells<br>DS stress: N = 4 animals, n = 14 cells | Two-way ANOVA | Interaction: F(1, 50) = 0.34, p = 0.56 |
| | | | | Genotype: F(1, 50) = 0.049, p = 0.83 |
| | | | | Stress: F(1, 50) = 19.29, p < 0.0001 |
| | | | Fisher's LSD post hoc test | WT control vs stress: p = 0.011 |
| | | | | DS control vs stress: p < 0.001 |
| 3D | sESPC rise time | WT control: N = 4 animals, n = 12 cells<br>WT stress: N = 6 animals, n = 15 cells<br>DS control: N = 4 animals, n = 14 cells<br>DS stress: N = 4 animals, n = 13 cells | Two-way ANOVA | Interaction: F(1, 50) = 0.83, p = 0.37 |
| | | | | Genotype: F(1, 50) = 0.056, p = 0.81 |
| | | | | Stress: F(1, 50) = 0.026, p = 0.87 |
| 3E | sEPSC decay time | WT control: N = 4 animals, n = 12 cells<br>WT stress: N = 6 animals, n = 15 cells<br>DS control: N = 4 animals, n = 13 cells<br>DS stress: N = 4 animals, n = 13 cells | Two-way ANOVA | Interaction: F(1, 49) = 0.74, p = 0.40 |
| | | | | Genotype: F(1, 49) = 2.40, p = 0.13 |
| | | | | Stress: F(1, 49) = 24.48, p < 0.0001 |
| | | | Fisher's LSD post hoc test | WT control vs stress: p = 0.005 |
| | | | | DS control vs stress: p < 0.001 |
| 3F | sEPSC AUC | WT control: N = 4 animals, n = 10 cells<br>WT stress: N = 6 animals, n = 14 cells<br>DS control: N = 4 animals, n = 13 cells<br>DS stress: N = 4 animals, n = 13 cells | Two-way ANOVA | Interaction: F(1, 46) = 0.92, p = 0.34 |
| | | | | Genotype: F(1, 46) = 7.20, p = 0.010 |
| | | | | Stress: F(1, 46) = 1.73, p = 0.20 |
| | | | Fisher's LSD post hoc test | Control WT vs DS: p = 0.24 |
| | | | | Stress WT vs DS: p = 0.011 |
| 3H | sIPSC amplitude | WT control: N = 4 animals, n = 12 cells<br>WT stress: N = 6 animals, n = 17 cells<br>DS control: N = 4 animals, n = 15 cells<br>DS stress: N = 4 animals, n = 16 cells | Two-way ANOVA | Interaction: F(1, 56) = 0.31, p = 0.58 |
| | | | | Genotype: F(1, 56) = 4.24, p = 0.044 |
| | | | | Stress: F(1, 56) = 7.12, p = 0.01 |
| | | | Fisher's LSD post hoc test | WT control vs stress: p = 0.15 |
| | | | | DS control vs stress: p = 0.023 |
| | | | | Control WT vs DS: p = 0.32 |
| | | | | Stress WT vs DS: p = 0.055 |
| 3I | sIPSC frequency | WT control: N = 4 animals, n = 13 cells<br>WT stress: N = 6 animals, n = 13 cells<br>DS control: N = 4 animals, n = 16 cells<br>DS stress: N = 4 animals, n = 15 cells | Two-way ANOVA | Interaction: F(1, 53) = 0.84, p = 0.36 |
| | | | | Genotype: F(1, 53) = 5.67, p = 0.021 |
| | | | | Stress: F(1, 53) = 1.98, p = 0.16 |
| | | | Fisher's LSD post hoc test | Control WT vs DS: p = 0.33 |
| | | | | Stress WT vs DS: p = 0.018 |
| 3J | sIPSC rise time | WT control: N = 4 animals, n = 13 cells<br>WT stress: N = 6 animals, n = 17 cells<br>DS control: N = 4 animals, n = 15 cells<br>DS stress: N = 4 animals, n = 16 cells | Two-way ANOVA | Interaction: F(1, 57) = 0.88, p = 0.35 |
| | | | | Genotype: F(1, 57) = 0.31, p = 0.58 |
| | | | | Stress: F(1, 57) = 4.26, p = 0.044 |
| | | | Fisher's LSD post hoc test | WT control vs stress: p = 0.041 |
| | | | | DS control vs stress: p = 0.42 |

| | | | | |
|---|---|---|---|---|
| 3K | sISPC decay time | WT control: N = 4 animals, n = 13 cells<br>WT stress: N = 6 animals, n = 17 cells<br>DS control: N = 4 animals, n = 15 cells<br>DS stress: N = 4 animals, n = 13 cells | Two-way ANOVA | Interaction: F(1, 54) = 4.19, p = 0.046 |
| | | | | Genotype: F(1, 54) = 0.43, p = 0.52 |
| | | | | Stress: F(1, 54) = 0.61, p = 0.44 |
| | | | Fisher's LSD post hoc test | WT control vs stress: p = 0.37 |
| | | | | DS control vs stress: p = 0.054 |
| | | | | Control WT vs DS: p = 0.34 |
| | | | | Stress WT vs DS: p = 0.058 |
| 3L | sIPSC AUC | WT control: N = 4 animals, n = 12 cells<br>WT stress: N = 6 animals, n = 17 cells<br>DS control: N = 4 animals, n = 14 cells<br>DS stress: N = 4 animals, n = 13 cells | Two-way ANOVA | Interaction: F(1, 52) = 0.66, p = 0.42 |
| | | | | Genotype: F(1, 52) = 0.43, p = 0.52 |
| | | | | Stress: F(1, 52) = 9.27, p = 0.004 |
| | | | Fisher's LSD post hoc test | WT control vs stress: p = 0.01 |
| | | | | DS control vs stress: p = 0.12 |
| 4B | % potentiation | WT control: n = 8<br>WT stress: n = 6<br>DS control: n = 8<br>DS stress: n = 7 | Two-way ANOVA | Interaction: F(1, 25) = 0.26, p = 0.62 |
| | | | | Genotype: F(1, 25) = 0.82, p = 0.37 |
| | | | | Stress: F(1, 25) = 14.13, p < 0.001 |
| | | | Fisher's LSD post hoc test | WT control vs stress: p = 0.007 |
| | | | | DS control vs stress: p = 0.027 |
| 4C | LTP paired pulse ratio | WT control: n = 8<br>WT stress: n = 6<br>DS control: n = 8<br>DS stress: n = 7 | Two-way ANOVA | Interaction: F(1, 25) = 0.02, p = 0.89 |
| | | | | Genotype: F(1, 25) = 5.36, p = 0.03 |
| | | | | Stress: F(1, 25) = 0.96, p = 0.34 |
| | | | Fisher's LSD post hoc test | Control WT vs DS: p = 0.08 |
| | | | | Stress WT vs DS: p = 0.16 |
| 4F | Rectification Index | WT control: N = 4 animals, n = 15 cells<br>WT stress: N = 4 animals, n = 14 cells<br>DS control: N = 4 animals, n = 15 cells<br>DS stress: N = 4 animals, n = 16 cells | Two-way ANOVA | Interaction: F(1, 56) = 1.58, p = 0.21 |
| | | | | Genotype: F(1, 56) = 1.15, p = 0.29 |
| | | | | Stress: F(1, 56) = 6.54, p = 0.013 |
| | | | Fisher's LSD post hoc test | WT control vs stress: p = 0.01 |
| | | | | DS control vs stress: p = 0.35 |
| 4G | AMPA/NMDA ratio | WT control: N = 4 animals, n = 15 cells<br>WT stress: N = 4 animals, n = 12 cells<br>DS control: N = 4 animals, n = 16 cells<br>DS stress: N = 4 animals, n = 12 cells | Two-way ANOVA | Interaction: F(1, 51) = 4.16, p = 0.046 |
| | | | | Genotype: F(1, 51) = 6.95, p = 0.011 |
| | | | | Stress: F(1, 51) = 0.58, p = 0.45 |
| | | | Fisher's LSD post hoc test | WT control vs stress: p = 0.055 |
| | | | | DS control vs stress: p = 0.37 |
| | | | | Control WT vs DS: p = 0.65 |
| | | | | Stress WT vs DS: p = 0.003 |
| 4H | Paired Pulse ratio | WT control: N = 4 animals, n = 15 cells<br>WT stress: N = 4 animals, n = 14 cells<br>DS control: N = 4 animals, n = 15 cells<br>DS stress: N = 4 animals, n = 16 cells | Two-way ANOVA | Interaction: F(1, 56) = 1.21, p = 0.28 |
| | | | | Genotype: F(1, 56) = 0.18, p = 0.67 |
| | | | | Stress: F(1, 56) = 1.95, p = 0.17 |

| | | | | |
|---|---|---|---|---|
| 4I | NMDA EPSC decay time | WT control: N = 4 animals, n = 16 cells<br>WT stress: N = 4 animals, n = 15 cells<br>DS control: N = 4 animals, n = 16 cells<br>DS stress: N = 4 animals, n = 16 cells | Two-way ANOVA | Interaction: F(1, 59) = 0.36, p = 0.55 |
| | | | | Genotype: F(1, 59) = 1.44, p = 0.23 |
| | | | | Stress: F(1, 59) = 2.12, p = 0.15 |
| 4J | AMPA EPSC decay time | WT control: N = 4 animals, n = 16 cells<br>WT stress: N = 4 animals, n = 14 cells<br>DS control: N = 4 animals, n = 15 cells<br>DS stress: N = 4 animals, n = 15 cells | Two-way ANOVA | Interaction: F(1, 56) = 0.002, p = 0.96 |
| | | | | Genotype: F(1, 56) = 1.75, p = 0.19 |
| | | | | Stress: F(1, 56) = 5.38, p = 0.024 |
| | | | Fisher's LSD post hoc test | WT control vs stress: p = 0.11 |
| | | | | DS control vs stress: p = 0.1 |
| 4K | PP AMPA EPSC decay time ratio | WT control: N = 4 animals, n = 16 cells<br>WT stress: N = 4 animals, n = 15 cells<br>DS control: N = 4 animals, n = 15 cells<br>DS stress: N = 4 animals, n = 15 cells | Two-way ANOVA | Interaction: F(1, 57) = 0.17, p = 0.68 |
| | | | | Genotype: F(1, 57) = 0.90, p = 0.35 |
| | | | | Stress: F(1, 57) = 0.003, p = 0.96 |
| 4L | NMDA EPSC AUC | WT control: N = 4 animals, n = 16 cells<br>WT stress: N = 4 animals, n = 15 cells<br>DS control: N = 4 animals, n = 13 cells<br>DS stress: N = 4 animals, n = 16 cells | Two-way ANOVA | Interaction: F(1, 56) = 15.20, p < 0.001 |
| | | | | Genotype: F(1, 56) = 1.64, p = 0.21 |
| | | | | Stress: F(1, 56) = 0.47, p = 0.50 |
| | | | Fisher's LSD post hoc test | WT control vs stress: p = 0.024 |
| | | | | DS control vs stress: p = 0.002 |
| | | | | Control WT vs DS: p = 0.075 |
| | | | | Stress WT vs DS: p < 0.001 |
| 4M | AMPA EPSC AUC | WT control: N = 4 animals, n = 16 cells<br>WT stress: N = 4 animals, n = 14 cells<br>DS control: N = 4 animals, n = 15 cells<br>DS stress: N = 4 animals, n = 15 cells | Two-way ANOVA | Interaction: F(1, 59) = 2.85, p = 0.10 |
| | | | | Genotype: F(1, 59) = 1.76, p = 0.19 |
| | | | | Stress: F(1, 59) = 9.17, p = 0.004 |
| | | | Fisher's LSD post hoc test | WT control vs stress: p = 0.35 |
| | | | | DS control vs stress: p = 0.001 |
| 4N | PP AMPA EPSC AUC ratio | WT control: N = 4 animals, n = 16 cells<br>WT stress: N = 4 animals, n = 15 cells<br>DS control: N = 4 animals, n = 16 cells<br>DS stress: N = 4 animals, n = 15 cells | Two-way ANOVA | Interaction: F(1, 58) = 0.007, p = 0.93 |
| | | | | Genotype: F(1, 58) = 0.09, p = 0.77 |
| | | | | Stress: F(1, 58) = 0.038, p = 0.85 |
| T1A | Membrane resistance | WT control: N = 5 animals, n = 33 cells<br>WT stress: N = 7 animals, n = 39 cells<br>DS control: N = 5 animals, n = 37 cells<br>DS stress: N = 7 animals, n = 33 cells | Two-way ANOVA | Interaction: F(1, 138) = 0.66, p = 0.42 |
| | | | | Genotype: F(1, 138) = 0.20, p = 0.65 |
| | | | | Stress: F(1, 138) = 4.54, p = 0.035 |
| | | | Fisher's LSD post hoc test | WT control vs stress: p = 0.35 |
| | | | | DS control vs stress: p = 0.041 |
| T1B | Cell capacitance | WT control: N = 5 animals, n = 34 cells<br>WT stress: N = 7 animals, n = 39 cells<br>DS control: N = 5 animals, n = 36 cells<br>DS stress: N = 7 animals, n = 26 cells | Two-way ANOVA | Interaction: F(1, 131) = 1.34, p = 0.25 |
| | | | | Genotype: F(1, 131) = 3.35, p = 0.07 |
| | | | | Stress: F(1, 131) = 14.96, p < 0.001 |
| | | | Fisher's LSD post hoc test | WT control vs stress: p < 0.001 |
| | | | | DS control vs stress: p = 0.069 |
